# Intracellularly Coupled Oscillators for Synthetic Biology

**DOI:** 10.1101/2025.09.23.678067

**Authors:** Gábor Holló, Jung Hun Park, Rose Evard, Yolanda Schaerli

**Affiliations:** Department of Fundamental Microbiology, University of Lausanne, Biophore Building, 1015, Lausanne, Switzerland

## Abstract

Synthetic biology aims to engineer or re-engineer living systems. To achieve increasingly complex functionalities, it is beneficial to use higher-level building blocks. In this study, we focus on oscillators as such building blocks, propose novel oscillator-based circuit designs and model the interactions of intracellularly coupled oscillators. We classify these oscillators on the basis of coupling strength: independent, weakly or strongly, and deeply coupled. We predict a range of fascinating dynamic behaviours to arise in these systems, such as the beat phenomenon, amplitude and frequency modulation, period doubling, higher-period oscillations, chaos, resonance, and synchronization, with the aim of guiding future experimental work in bacterial synthetic biology. Finally, we outline potential applications, including oscillator-based computing that integrates processing and memory functions, offering multistate and nonlinear processing capabilities.

## Introduction

In synthetic biology (SynBio), the goal is to engineer living organisms to perform specific, useful tasks by designing and constructing biological systems [1–4]. SynBio is inherently multidisciplinary, integrating principles from fields such as molecular biology, genetics, electrical engineering, and computer science. Similar to electrical engineering, where simple components like resistors, transistors, and capacitors are assembled to create functional circuits, SynBio uses biological building blocks, such as genes, proteins, and regulatory elements, to construct biological “circuits” that control cellular behavior in predictable ways. As electrical circuits become more complex, higher-level components such as microchips are often employed to streamline the design process. Similarly, in SynBio, achieving more advanced and intricate functionalities could benefit from using higher-level biological building blocks, such as multiple oscillators coupled to one another, which can produce complex, diverse, and useful behaviors.

Insights from physics, chemistry, biology, and engineering show that coupled oscillators exhibit remarkable dynamic behaviors. These phenomena have revolutionized industries over the past centuries, with applications in telecommunications [5], micro-electromechanical systems (MEMS) [6], healthcare (e.g. cardiac pacemakers) [7], and quantum computing [8], among others. Beyond practical applications, the study of coupled oscillators enhances our understanding of natural processes such as synchronization in circadian rhythms [9], neuronal activity [10], mechanical systems [11], and even animal and human behaviors [12–14].

In biological systems, natural genetic oscillators are coupled at both intracellular and intercellular levels. For instance, cell division and the circadian clock are intracellularly coupled, enabling them to achieve robust synchronization [15, 16]. Similarly, metabolic oscillations have been shown to synchronize with the early and late stages of the cell cycle, as demonstrated in a study in budding yeast [17]. Intercellular coupling also plays a pivotal role in processes like presomitic mesoderm segmentation during vertebrate embryonic development. Precise spatiotemporal oscillations are essential for the proper formation of vertebrae [18–20]. Another example of coupling can be observed in plant growth, whereby the growth rate and shoot curvature oscillate in anti-phase of one another, effectively dampening curvature and enabling straight growth [21, 22]. These examples highlight the critical role of coupled oscillations in maintaining coordination and functionality across biological systems.

In this study, we explore the potential phenomena that may arise when multiple synthetic oscillatory circuits are introduced within a single living cell, e.g., intracellular coupling. Our goal is to contribute to fundamental advancements in SynBio, just as coupled oscillators have done in many other fields. Figure 1 gives an overview of the oscillators we use in this study as building blocks for constructing coupled oscillatory systems. The first oscillator, the Goodwin oscillator [23–26] consists of a single self-repressing node and exhibits fast, low-amplitude oscillations. As the time period increases, the amplitude likewise grows until protein levels approach saturation. The dual-feedback oscillator, introduced by Stricker *et al.* [26], is the second fastest oscillator in this lineup. The repressilator, designed by Elowitz *et al.* in 2000 [27], was the first synthetic oscillator in synthetic biology. Since then, numerous variants and alternative circuit topologies have been developed, spanning a wide range of time periods [26, 28–33]. While these designs predominantly rely on transcription factor-mediated regulation, the CRISPRlator, our slowest oscillator, relies on CRISPR interference-based repression [31]. These oscillators span a broad range of amplitudes and time periods, enabling the study of fascinating behaviours and paving the way for innovative applications in the future.

**Figure 1:**
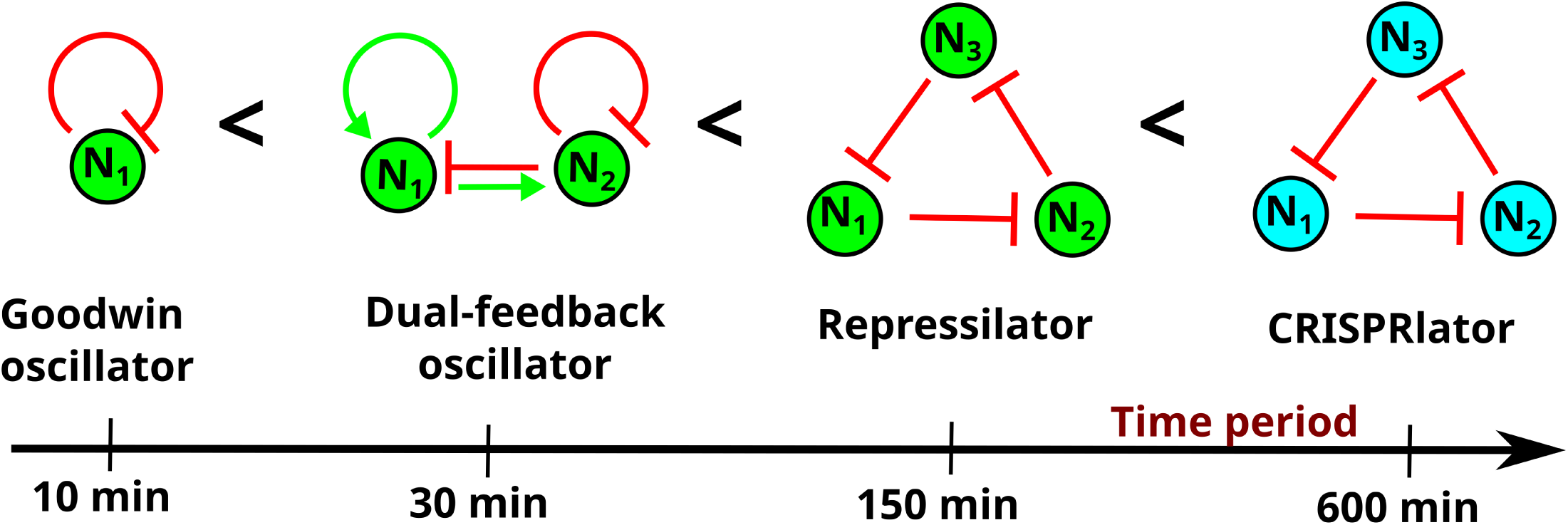
Oscillators used in this study. The time period increases from left to right. While representative time periods are shown, these can vary significantly depending on the parameters used in simulations, biological implementation, or experimental conditions. The blue nodes in the CRISPRlator symbolize that this circuit relies on CRISPR-based repression rather than transcription factor interactions.

One possibility to study coupled oscillators is to use *non-autonomous* systems and investigate the interactions between an oscillator inside the cell and an external oscillatory input [33–35]. For example, we previously examined how oscillatory light inputs interact with an optogenetic repressilator, referred to as the “optoscillator”. [33]. On the other hand, in autonomous systems, each oscillator operates based on its own internal dynamics, and interactions occur between oscillators, without external time-dependent forcing. Several studies have explored synthetic intercellularly coupled oscillators mediated by small-molecule-based cell-cell communication [28, 29, 36–39]. These efforts focused on populations in which every cell harbors an identical oscillator. Such systems have proven to be a powerful tool for constructing complex systems and investigating intricate dynamic behaviors [28, 29, 36–38].

However, intercellular coupling introduces inherent limitations: signaling molecules are required to traverse cell membranes, diffuse through the extracellular space and re-enter neighboring cells in order to influence other oscillators [39]. This process can significantly attenuate and delay signal transmission. Moreover, studying interactions between distinct oscillators in such setups requires the use of different cell types and the establishment of stable co-cultures, posing additional experimental challenges [40]. In contrast, intracellularly coupled oscillators enable alternative coupling strategies that bypass these delays and signal degradation, offering a more direct and efficient framework for exploring oscillator interactions.

Notably, intracellular coupling between oscillators can be achieved through direct regulatory interactions, in which components of one circuit modulate the dynamics of another. In addition to these explicit connections, it is well established that competition for shared, limited cellular resources, such as RNA polymerases, ribosomes, proteases, dCas9 and nutrients, can influence the behavior of otherwise orthogonal genetic circuits [41–48]. While such competitions are typically considered undesirable, they can be deliberately harnessed to couple synthetic circuits. For instance, Prindle and colleagues demonstrated that protease competition can be harnessed to couple oscillators [44]. A similar mechanism can be envisioned for CRISPRi-based systems, where competition for dCas9 could serve as a coupling strategy. Metabolic networks offer an additional layer of interaction between genetic circuits. Oscillations in metabolic activity can naturally synchronize distinct circuits via fluctuations in shared metabolites [49]. For example, changes in substrate or product availability can modulate oscillator period and amplitude [50]. A notable case is the metabolator, a synthetic metabolic oscillator driven by dynamic flux between interconverting metabolite pools [51]. Thus, intracellular oscillator coupling presents a range of intriguing possibilities, which we explore in this study.

While coupled oscillators have been extensively studied in physics and engineering, their systematic design within single living cells remains largely unexplored. Previous efforts in synthetic biology have focused mainly on externally driven or intercellularly coupled systems [26, 28, 29, 33–35, 37, 39]. Here, we introduce a framework for autonomous intracellular coupling between synthetic oscillators, enabling complex dynamical behaviors, such as synchronization, resonance, and chaos, to emerge solely from internal regulatory interactions. By treating oscillators as modular building blocks, this work establishes a new design paradigm that bridges dynamical systems theory with the practical challenges of scalable genetic circuit engineering.

Figure 2 provides an overview of the classification of coupled oscillators relevant to this study. Coupled oscillators can be divided into two main categories: *non-autonomous* and *autonomous* systems. Autonomous systems are self-governing and time-invariant, described by differential equations in which the independent variable (time) does not explicitly appear:

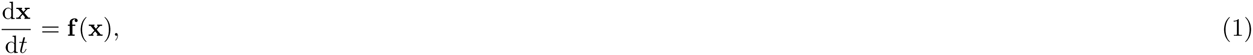

**Figure 2:**
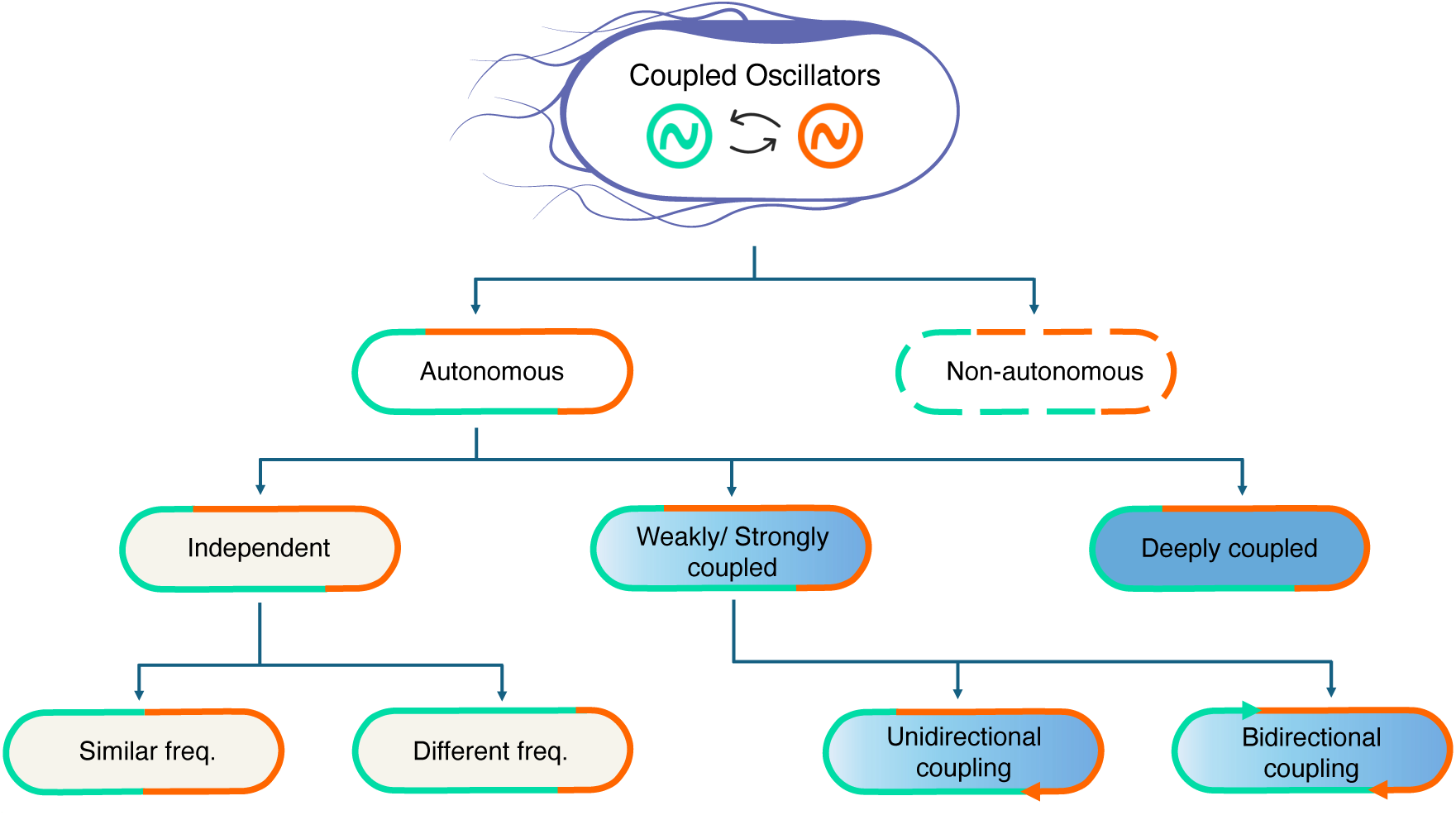
Overview of the classification of intracellular coupled oscillators relevant to this study. In non-autonomous systems, oscillators within cells are influenced by an external force (dashed orange and turquoise lines). In autonomous systems, however, the oscillators interact directly with each other within the cells (solid orange and turquoise lines). Depending on these interactions, different category of oscillators may emerge. Independent oscillators occur when no interactions are present (light beige background), while deeply coupled oscillators (dark blue background) arise under infinitely strong interactions. Between these extremes, oscillators can be classified as weakly or strongly coupled (gradient of blue intensity background). Further distinctions can be made regarding the relative frequency of the oscillators (length of orange/turquoise lines) or the direction of coupling (orange or turquoise arrows).

where **x** denotes the state vector of the system and **f** (**x**) defines the intrinsic dynamics. In contrast, non-autonomous systems depend explicitly on time, either through external driving forces or time-varying parameters:

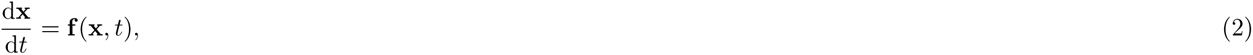

where the additional argument *t* captures the influence of time-dependent inputs or modulations. This distinction is fundamental in classifying oscillatory dynamics, particularly in biological contexts where intrinsic rhythms (autonomous) can be perturbed or entrained by external cues such as light–dark cycles or chemical stimuli (non-autonomous).

This study focuses on autonomous coupled oscillators, where multiple oscillators interact without external influences. However, as we will show, autonomous unidirectionally coupled systems share notable similarities with their non-autonomous counterparts. We further classify autonomous oscillators on the basis of the strength of their interactions. When the interaction is negligible, we refer to them as *independent* oscillators, whereas with infinitely strong interaction, we observe *deeply coupled* oscillators. Between these extremes are *weakly* and *strongly coupled oscillators*. The direction of coupling interaction can be taken into account when categorizing coupling oscillators as well. When one oscillator influences the other without being affected in return (or the effect is negligible), this interaction is referred to as unidirectional or master-slave coupling. In contrast, when both oscillators mutually influence each other, it is referred to as bidirectional or peer-to-peer coupling [52–54]. Another critical aspect influencing the behavior of coupled oscillators is their relative frequencies.

Since we are utilizing higher-level building blocks (i.e., oscillators) to create these coupled oscillators, establishing simulations at a higher level of abstraction is straightforward. For this purpose, we used the GRN modeler [55], our previously developed tool to simulate, and analyze gene regulatory networks (GRNs). It features a user-friendly graphical user interface (GUI) that allows us to construct and model the circuits node by node, significantly accelerating the design and modeling of GRNs.

For our simulations, we employed three distinct models. The first model, developed by Elowitz *et al.* [27], represents the repressilator. In other cases, such as when describing the Goodwin oscillator [23], a more detailed model is required to account for the role of enzymatic protein degradation. For these situations, we employed the model created by Tomazou *et al.* [56]. For the CRISPRlator, we used a model described by Holĺo *et al.* [55]. We provided a comprehensive summary of these models in our previous work [55], and for the reader’s convenience, we summarize them again in the SI Section 1. Throughout this study, we will refer to these models as the Elowitz, Tomazou and CRISPR models, respectively.

One promising application of coupled oscillators is their use in oscillator-based computing (OBC), enabling information processing in response to inputs, an approach also referred to as biocomputing. Most current biocomputing implementations in synthetic biology rely on the logic-gate metaphor from electronics, producing binary outputs [57, 58]. However, this framework is inherently limiting for complex computations and may not represent the most suitable paradigm for information processing in living cells [59–61]. In OBCs, the information is encoded in the frequency or phase of the oscillators, rather than in the signal’s strength, mimicking the natural neuronal oscillations of the brain [62]. In the brain, oscillations couple with one another in phase and/or frequency to coordinate complex calculations/actions such as visual processing, memory, cognition, and voluntary movement generation [10, 63–65]. OBCs aim to reproduce this behavior, with individual processing units (“neurons”) performing basic calculations in response to multiple signals [66]. The interactions between oscillators then allow complex, nonlinear calculation functions as the output from the collection of neurons (see Csaba & Porod (2020) [66] for a comprehensive review about OBCs). While researchers in other fields have attempted to emulate the functioning of the brain’s action in biological systems [67–69], to the best of our knowledge, OBCs have not yet been designed in SynBio.

Here, we computationally investigate how the topology and coupling strength of intracellular oscillators influence their behavior, offering new insights into the dynamics of intracellular systems. This theoretical framework lays a foundation for experimental efforts to engineer novel oscillatory circuits. After exploring different scenarios that can arise with intracellularly coupled oscillators, we study their implementation in OBCs and their potential application in SynBio. By presenting this foundational concept, we aim to inspire further research and innovation in the field.

## Results

### Independent oscillators

We began by examining the simplest case of autonomous coupled oscillators, two systems operating without direct interactions. This scenario involves two independent oscillators controlling the state of a common node through logic gates. Although the assumption of complete independence is highly idealized, as in real cells, oscillators often share molecular components or are interconnected through metabolic activities, it remains valuable to study these systems as a limiting case.

Using the Tomazou model [56], we simulated two independently coupled repressilators with similar frequencies (Figure 3I). We kept the parameters of the second oscillator constant while gradually varying the concentration of the PROT_1_ protease, which degrades the proteins of the first repressilator. Although the protease concentration was held constant in the simulations, its activity varied dynamically due to oscillations in the concentrations of its substrates (Figure S6). The adjustment of the protease concentration allowed us to alter the degradation rate of the transcription factors in the first repressilator, thereby modifying its oscillation period, while the period of the second remains constant. Both oscillators control the output node through a NOR gate, resulting in an output signal only when both oscillators are in-phase, i.e., when N_1_ and N_4_ are deactivated simultaneously. Due to the slight frequency difference between the oscillators, the phase difference is changing continuously between them, leading to periodic activation and deactivation of the output node. This interference pattern, known from acoustics as the *beat phenomenon*, can be calculated with the following formula [70]:

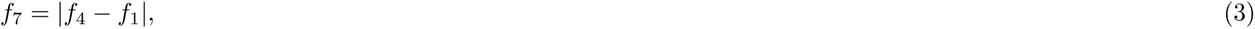

where *f*_4_ and *f*_1_ are the frequencies of the independent repressilators, and *f*_7_ is the beat frequency at the output node (N7). According to this formula, when the frequency difference between the two oscillators is small, the resulting beat frequency is also low, leading to a prominent peak at a long beat period (Figure 3I,d). As shown in this figure and determined by Eq. 3, the time period of the output node, calculated from trajectories with varying protease concentrations, corresponds closely to the period of the beat phenomenon. This ideal case allows us to easily calculate the necessary protease concentration for a given output period.

**Figure 3:**
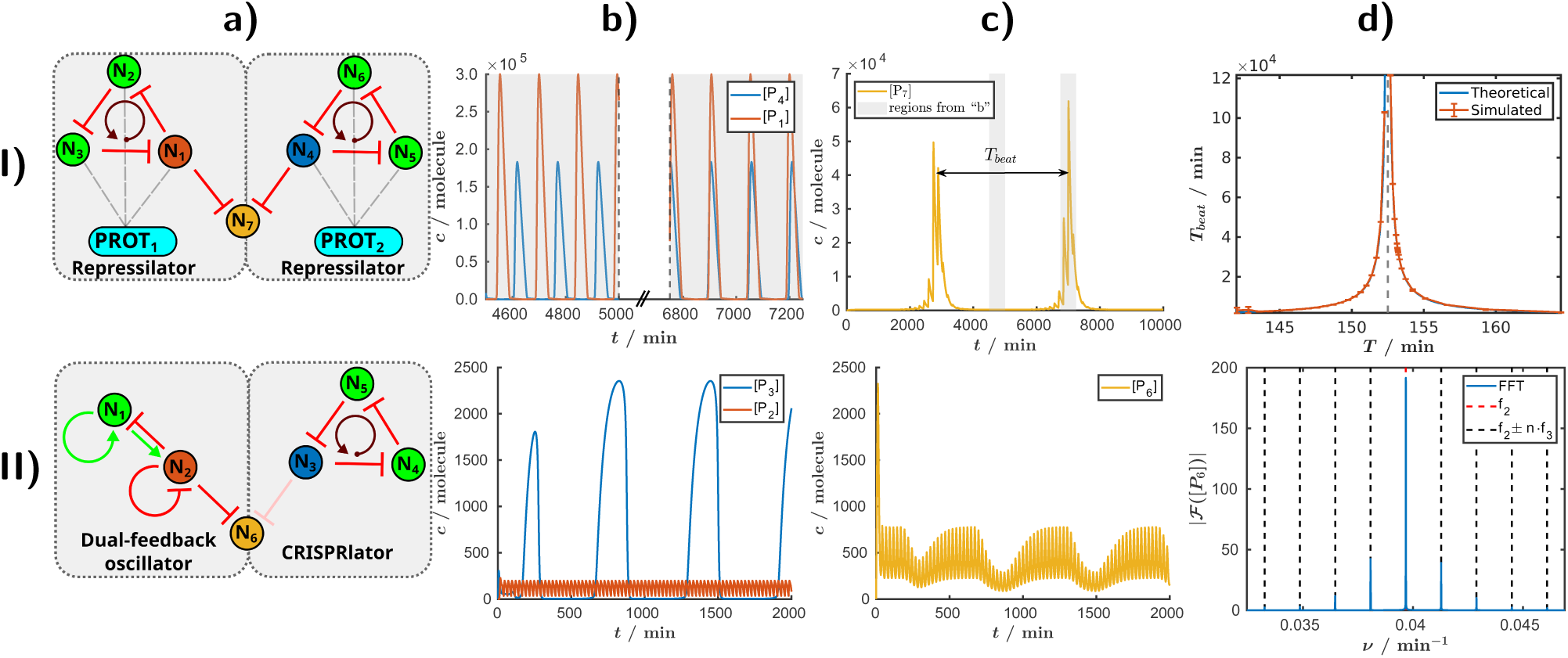
Independently Coupled Oscillators. Row I: The Beat Phenomenon. **a)** Circuit topology of two repressilators coupled via a NOR gate at node N_7_. **b)** Protein concentration trajectories of nodes N_1_ (red) and N_4_ (blue), respectively producing proteins P_1_ and P_4_. Two segments of the trajectories are presented: one up to 5000min, illustrating distinct peak separation, and another from 6800 min onward, where the peaks overlap. c) Oscillations in the output node N_7_. Peaks are observed when the oscillations in (b) are in-phase (second half of the graph). Conversely, when the oscillations in (b) are out-of-phase (first half), N_7_ remains repressed. Gray rectangles indicate the time ranges corresponding to panel b. d) Comparison of theoretical (blue) and simulated (red) beat time period as a function of N_1_’s oscillation period. **Row II: Amplitude Modulation**. **a)** Circuit topology of a dual-feedback oscillator coupled to a CRISPRlator oscillator via N_6_. **b)** Protein concentration dynamics of nodes N_2_ (red) and N_3_ (blue). c) Output node N_6_ oscillations. d) Fourier transform (FFT) of N_6_ (blue), with expected modulation frequency peaks (black dashed lines). *f*_2_ and *f*_3_ denote the frequencies of N_2_ (dual-feedback oscillator) and N_3_ (CRISPRlator), respectively. The detailed descriptions of the models are available in “beat.html” and in “AM independent Stricker weak.html”, respectively. An extended figure caption with additional details is available in SI Section 2.

For practical applications, this method provides a way to achieve significantly different frequencies without redesigning the oscillatory circuits. Typically, circuit adjustments to alter its frequency (e.g., by modifying promoter strength or protease concentration) only tune the frequency within a limited range. Significant changes often require the design of a new circuit. Leveraging the beat phenomenon, however, allows substantial adjustments to the output node’s period without the need for new circuit designs, making it a valuable technique for tuning oscillatory systems.

Next, we examined a scenario involving independently coupled autonomous oscillators with significantly different time periods. We coupled a CRISPRlator [31, 55, 71] and a dual-feedback oscillator [26] using again a NOR gate (Figure 3II,a). In our model, the time period of the CRISPRlator is 604.5 minutes, while the time period of the dual-feedback oscillator with the modified Elowitz model is 25.2 minutes (Figure 3II,b) (see the extended figure caption in SI Section 2 for details of the model). We observed a phenomenon known as *amplitude modulation* (AM), in which the amplitude of a high-frequency carrier signal is modulated by a lower-frequency signal. Specifically, the carrier signal from the fast dual-feedback oscillator (P_2_, red) is modulated by the slow CRISPRlator (P_3_, blue), resulting in an AM signal at the output node (P_6_, yellow) (Figure 3II,b,c). In this simulation, the repression strength from N_3_ to N_6_ is low (0.01%) ensuring a continuous carrier signal. For comparison, the scenario with complete repression is shown in Figure S5. Figure 3II,d illustrates that under this ideal conditions with completely independent oscillators, the frequency of the output node is indeed determined by the modulation of the dual-feedback oscillator’s frequency, occurring at integer multiples of the CRISPRlator’s frequency. Furthermore, we confirm the independence of both frequency and phase between the uncoupled oscillators. The time-resolved frequency spectrum of the output signal is characterized using wavelet transformation (Figure S8) and the Hilbert transform (Figure S9), providing a detailed view of the modulation dynamics. Finally, we show that AM can also be achieved by coupling a repressilator and a CRISPRlator (Figure S7).

The basic principles underlying independently coupled oscillators, such as beat phenomenon and amplitude modulation, can be illustrated analytically using simple harmonic functions (see SI, Section 2).

### Deeply coupled oscillators

In contrast to independent oscillators, where the coupling strength is zero, in deeply coupled oscillators the coupling is infinitely strong. Although these circuits could be viewed as a single unified system rather than as coupled oscillators, treating them as coupled systems provides a more intuitive framework for understanding their behavior.

To study the behavior of deeply coupled oscillators, we connected two repressilators through a common edge (a repression) (Figure 4a), creating a scenario of deep coupling in which the oscillators cannot oscillate at different frequencies or phases. The “first repressilator” comprises the nodes N_1_, N_2_, and N_3_, while the “second repressilator” consists of N_1_, N_4_, and N_3_. We gradually adjusted the promoter strength of the fourth node (Figures 4b and c) – a node not connected to the common edge. As the promoter strength decreases, the dynamic behavior and time period of the circuit increasingly resemble those of a single repressilator. Initially, due to the coupling, the system exhibits asymmetry: nodes display varying amplitudes, and the overall period is longer than that of the standard repressilator (Figure 4b, first row). However, as the promoter strength of N_4_ decreases and the influence of the second oscillator weakens, the system becomes symmetrical (Figure 4b, third row), with its period converging to that of the three-node repressilator. We would like to emphasize that, in most cases, we highlight these phenomena in coupled oscillators using a single model and a specific example (e.g., in this case, by modifying the promoter strength of N_4_). However, the system’s oscillation frequency can be fine-tuned through multiple approaches, all of which produce qualitatively similar results to the one presented. For instance, frequency adjustments can be achieved by altering the promoter strength (Figure 4), modifying the protease concentration (Figure S11), or varying repression strength with a small chemical (Figure S12). Therefore, while this study outlines potential scenarios, experimental implementation will require selecting the most appropriate approach and modeling the corresponding conditions.

**Figure 4:**
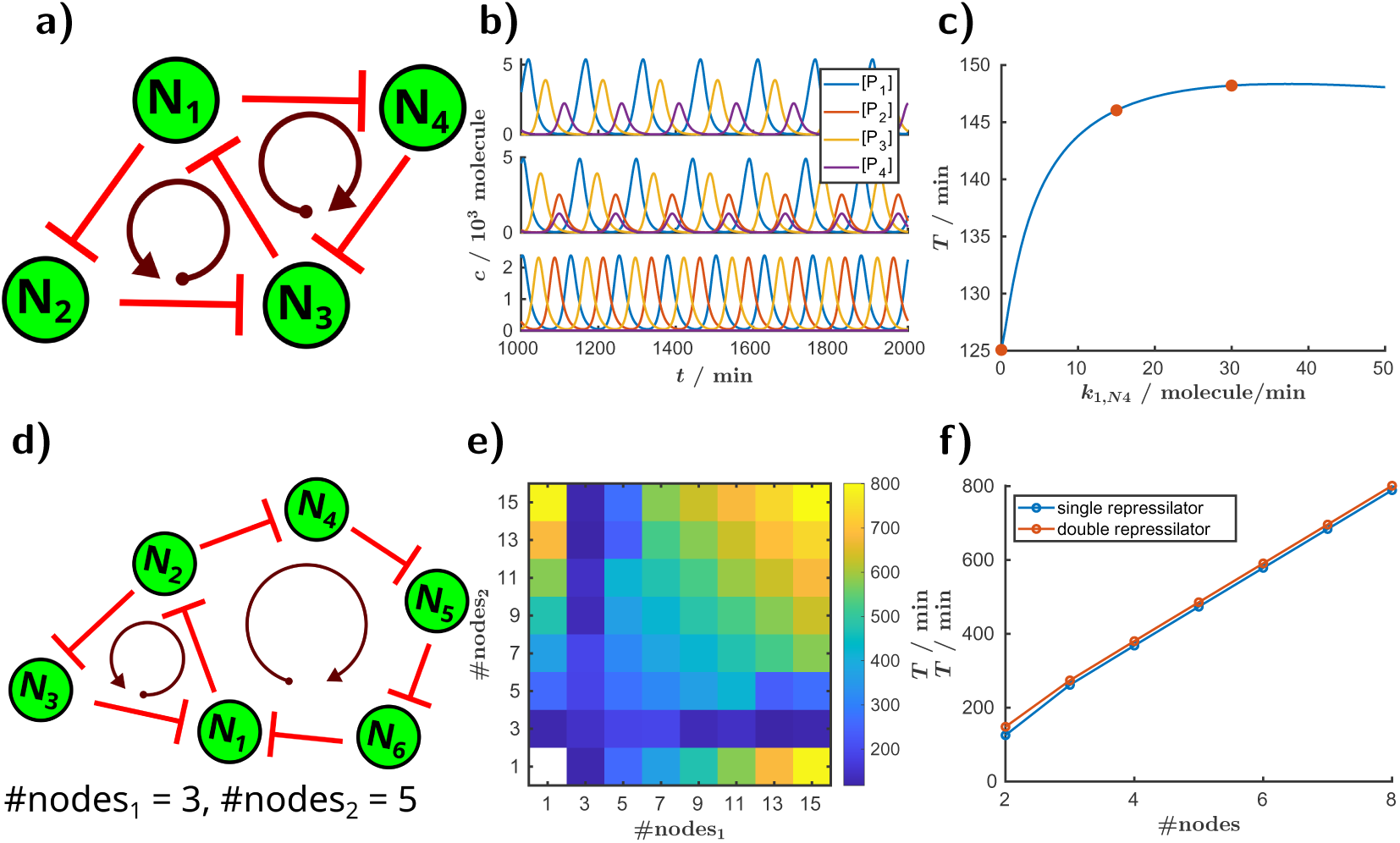
Deeply coupled repressilators with a shared edge. **a)** Topology of two deeply coupled three-node repressilators. **b)** Protein trajectories of corresponding nodes, with varying promoter strengths for the N_4_ node: 30, 15 and 0 molecule/minute, respectively. **c)** Time period of the coupled system as a function of the promoter strength of the fourth node. Scenarios from panel (b) are marked with red dots. **d)** Topology of deeply coupled repressilators with three and five nodes. **e)** Time period of the system as a function of the number of nodes in the coupled oscillators. **f)** Time period of a single repressilator and a two deeply coupledrepressilators sharing an edge with both rings having an equal number of nodes (”double repressilator”), plotted as a function of the number of nodes in a single ring. The simulations ran for 10^4^ minutes, with the first 10^3^ minutes regarded as transient time. The detailed descriptions of the model for the circuit in a) is available in “deeply coupled Elowitz.html”.

Next, we investigated the behavior of two deeply coupled oscillators when their frequencies vary significantly. We previously demonstrated that the time period of the repressilator family increases linearly with the number of nodes [55]. In Figures 4d and 4e, we couple two repressilators with odd numbers of nodes. In the matrix of Figure 4e, the two axes represent the number of nodes in the first and second circuits. A node number of 1 indicates the absence of the circuit. Therefore, the first column and the last row show the time period of the repressilator without coupling to a second oscillator, while the point (1, 1) represents a system without oscillators. The matrix is symmetric because the system itself is symmetric: the first and second oscillators are interchangeable. As expected, in the uncoupled case, the time period increases linearly with the number of nodes red(Figure S10). Interestingly, a similar linear relationship is observed when the two oscillators share an edge and have the same number of nodes, although the overall period is slightly longer due to the shared edge (Figure 4f). However, when a large circuit (such as a five- or fifteen-node system) is coupled via a shared edge to a smaller (and faster) oscillator, the overall system accelerates to the frequency of the faster oscillator. This occurs because along the faster route, the first common node is repressed earlier, forcing the frequencies and phases of the coupled oscillators to synchronize. As a potential application, this coupling strategy could be used to accelerate larger or slower oscillators by linking them to smaller and faster oscillators. The repressilator can only oscillate with an odd number of nodes; therefore, both of the coupled circuits in this case were designed with an odd number of nodes. If either circuit contains an even number of nodes, the system becomes bistable, and oscillations cease. However, a similar approach can be applied to circuits like the reptolator or actolator, which resemble the repressilator yet are capable of oscillating with an even number of nodes [55].

#### The deeply coupled Goodwin oscillator

Oscillatory circuits are often constructed from delayed negative feedback and self-activation motifs [23, 72]. The Goodwin oscillator represents one of the simplest examples, relying on a single self-repressing node that generates high-frequency, low-amplitude oscillations. We found that this characteristic “Goodwin-type” behavior is not restricted to a single node: it can also emerge in multi-node systems when the nodes become synchronized. For instance, the classical toggle switch [73, 74], typically known for bistable switching, can exhibit fast, low-amplitude oscillations when both nodes oscillate in phase. In this regime, the circuit behaves effectively as a single self-repressing unit.

Interestingly, even the repressilator, a canonical model of sequential (out-of-phase) oscillations, can transition to a Goodwin-type, in-phase oscillatory state under specific coupling conditions. The coexistence and interaction of these two oscillatory modes leads to rich dynamics, including bistability, period-doubling, and even chaotic behavior at strong coupling (Figure S13). These findings highlight that deeply coupled circuits can display unexpectedly diverse dynamical regimes, extending the concept of the Goodwin oscillator far beyond its traditional single-node form. A detailed analysis and supporting simulations are provided in SI Section 3.2.

### Weakly and strongly coupled oscillators

Previously we discussed the independent case, where oscillators do not interact, and the case of deep coupling, where the system behaves like a single oscillator. Next, we explored the cases that fall between these two extremes and that we further categorized into weakly and strongly coupled oscillators. Typically, most studies about coupled oscillators analyze these intermediate categories. Although there is no strict boundary or universally agreed-upon definition to distinguish weak from strong coupling, a common criterion for strong coupling is when the coupling strength (or energy exchange rate) exceeds the system’s loss rates [75, 76].

Independent oscillators exhibit no synchronization, while deeply coupled oscillators are fully synchronized. In contrast, weakly and strongly coupled oscillators display more nuanced synchronization dynamics, such as partial or limited synchronization. We first studied the effect of synchronization using a model of two coupled repressilators, based on the Tomazou model [56] (Figure 5a-d). Each oscillator is degraded by its own protease, but they also share a common protease, PROT_3_, allowing us to fine-tune the coupling strength by adjusting the concentration of this shared enzyme (Figure 5a). In Figure 5b, both oscillators have identical parameter sets, resulting in the same oscillation frequency. However, a small difference in the initial conditions leads to a phase offset between the oscillations. As we increase the coupling strength, represented by the number of protease molecules PROT_3_ (from 0 to 16), we observe the transition from independent oscillators (at zero coupling) where the phase difference remains constant, to synchronized oscillators where the phase difference diminishes. As expected, the stronger the coupling strength, the faster the oscillators synchronize.

**Figure 5:**
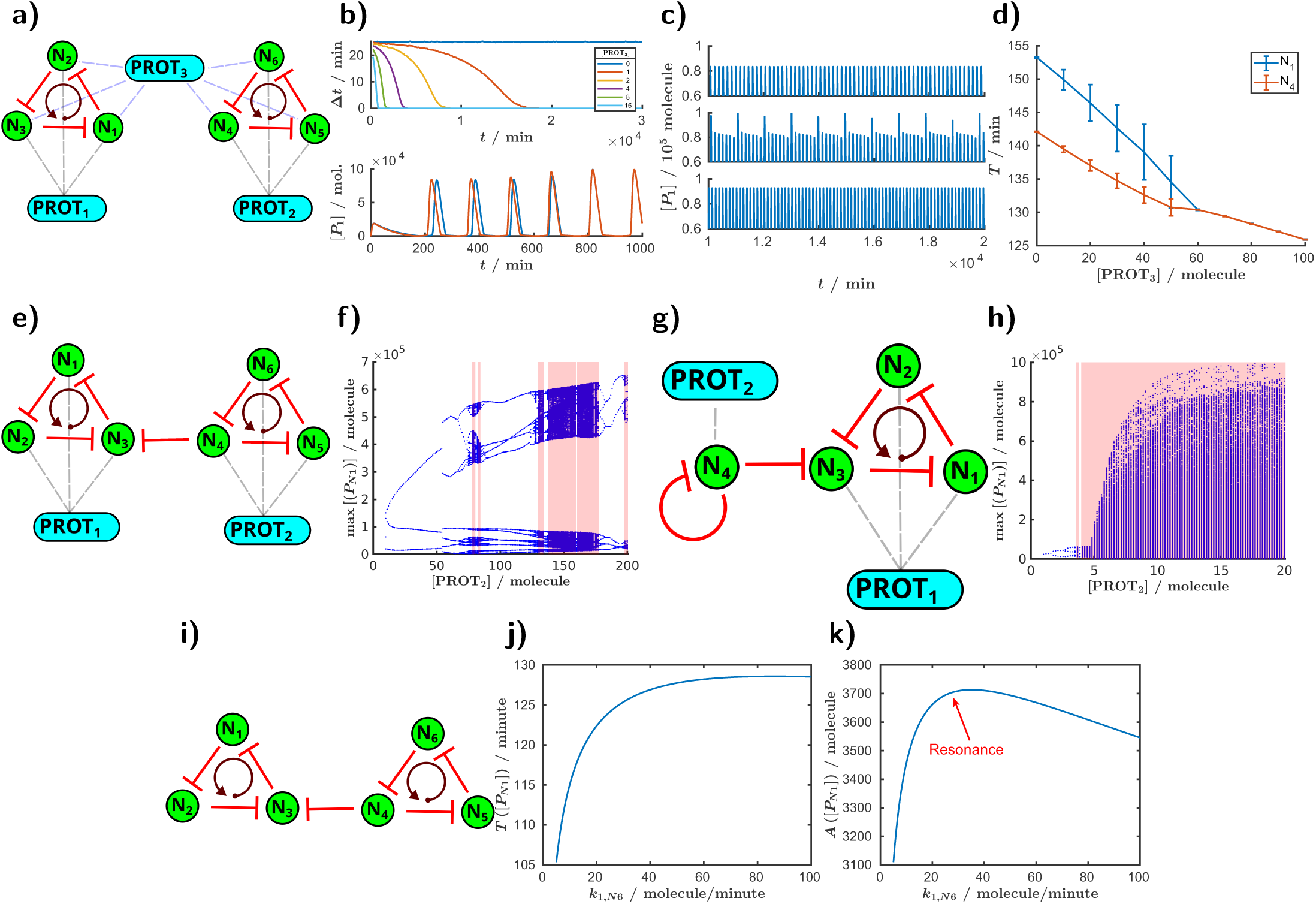
Synchronization, frequency and amplitude modulation, chaos, and resonance with coupled oscillators. a)-d) Synchronization in coupled repressilators. **a)** Topology of two oscillators coupled through PROT_3_. **b)** Phase difference and protein oscillations for varying PROT_3_ concentrations. **c)** Protein trajectories in N_1_ for increasing PROT_3_ levels (0, 40, and 100 molecules). **d)** Variation in average period for nodes 1 and 4 as a function of PROT_3_. **e)-h) Higher period oscillations and chaos. e)** Topology of two unidirectionally coupled repressilators. **f)** Bifurcation diagram for the system in (e), chaotic regions are depicted in red. **g)** Topology of a Goodwin oscillator coupled to a repressilator. **h)** Bifurcation diagram for the system in (g), chaotic regions are depicted in red. **i)-k) Resonance. i)** Topology of two unidirectionally coupled repressilators. **j)** Time period changes in the second oscillator with promoter strength variation in node N_6_. **k)** Oscillation amplitude at N_1_ with changes in the promoter strength of N_6_. Simulations for resonance (i-k) used the Elowitz model; others (a-h) used the Tomazou model. The details of the simulations can be found in “coupled repressilators Tomazou.html”, “repressilator repressilator.html”, “repressilator goodwin.html” and in “resonance.html”, respectively. An extended figure caption with additional details is available in SI Section 4

We then varied the protease concentrations in the coupled repressilators (PROT_1_ and PROT_2_), leading to differences in their natural frequencies (Figures 5c and 5d). With zero PROT_3_ concentration, representing the independent case, the oscillators exhibit regular oscillations (Figure 5c, top row), and their distinct time periods are evident at the start of the curves (Figure 5d). As the coupling strength increases, we observe higher-period oscillations with alternating large and small peaks (Figure 5c, middle row). This pattern arises from constructive and destructive interference, similar to the beat phenomenon seen in independent oscillators. However, as the coupling strength further increases, the oscillators’ time periods gradually converge, making the dynamics more complex compared to the uncoupled case. The error bars in Figure 5d indicate that the time period is not constant during the oscillation. When a critical coupling strength is reached, a bifurcation occurs, causing the individual frequencies of the oscillators to merge, resulting in a single, common time period. Beyond this critical point, the system achieves regular oscillations with constant amplitude and a stable time period (Figure 5c, bottom row). High phase-locking values (Figure S17I,c) indicate sustained synchronization between the oscillators, while the frequency perturbations observed in the continuous wavelet transform reflect dynamic adjustments in frequency during the synchronization process (Figure S17I). The instantaneous phase difference, calculated using the Hilbert transform (SI Section 2), varies during the synchronization process and stabilizes once the oscillators achieve full synchronization (Figure S17II).

For synchronization to occur, bidirectional coupling is typically required, whereas unidirectional coupling behaves similarly to non-autonomous systems. Previously, we experimentally demonstrated that non-autonomous systems can exhibit complex dynamics such as resonance, period-*N* oscillations or chaos [33]. A unidirectionally coupled system operates similarly, but with the periodic driving force coming from within the system rather than from an external source, suggesting that similar dynamic phenomena can be observed in these autonomous systems as well.

We already discussed amplitude modulation (AM) with independent coupled oscillators, where the oscillation frequency remains constant, and only the amplitude changes (Figure 3II). By unidirectionally coupling a slow oscillator to a faster oscillator, both FM (frequency modulation) and AM can be observed (Figure S18). As altering the promoter strength affects both amplitude and frequency, this generally results in imperfect FM and AM. However, using more complex topologies and building on previous studies [56, 77], it is possible to generate pure AM and FM signals with unidirectionally coupled oscillators (Figure S19).

We next explored the capability of unidirectionally coupled oscillators to produce chaotic oscillations, using two examples. In the first examples, a repressilator is driven by another repressilator (Figures 5e, f). and the frequency of the driving system is varied by adjusting the protease concentration, PROT_2_. For the coupled repressilators, we observe wide regions of higher-period oscillations and chaos (Figure 5e, f, and Figure S20).

In the second example, a Goodwin oscillator drives the repressilator (Figure 5g). Also here, the frequency of the Goodwin oscillator can be varied with the protease concentration. The observed chaotic region is notably wide (Figure 5h and Figure S20). This coupled oscillator presents a promising candidate for a synthetic chaotic autonomous oscillators. The large Lyapunov exponents facilitate the observation of chaotic dynamics experimentally, and the wide chaotic region reduces the need for precise experimental fine-tuning. Moreover, the system is relatively simple, consisting of only four nodes, which minimizes the potential burden on cells.

Apart from chaotic oscillations, resonance is another phenomenon of interest in coupled oscillators. In our previous work [33], we demonstrated both experimentally and computationally that the non-autonomous optoscillator can exhibit resonance (alongside higher-period oscillations and chaos). Our simulations were based on the Elowitz model [27]. Here we predict that resonance can also occur in autonomous coupled oscillators by using two unidirectionally coupled repressilators. In our simulations, we varied the frequency of the driving oscillator by changing the promoter strength while keeping the frequency of the other oscillator constant. By adjusting the promoter strength of the N_6_ node in the master oscillator, we found that the amplitude of protein oscillations achieves its maximum when this parameter is set around 30 molecules per minute (Figure 5o), which coincides with the point at which the two oscillators have equal frequencies.

### An Application: Oscillator-Based Computing

Finally, we explored an application of coupled oscillators in oscillator-based computing (OBC): the creation of information processing units with memory functions. To showcase this application, we studied two coupled CRISPRlators (Figure 6). The coupling is achieved through a shared dCas pool rather than a protease (as was the case in Figure 5). If the dCas concentration is high, there is no competition between the oscillators for the shared species, allowing each oscillator to function independently (Figure S23). In contrast, at low dCas concentrations, CRISPRlators cannot oscillate at different frequencies because a synchronized peak alignment would excessively deplete the dCas pool. This constraint enforces synchronization with a fixed phase difference between oscillators. It is important to note that these simulations only account for the effects related to the dCas pool. In a real system, higher synchronization could arise from factors such as proteases or cellular metabolic activity, which would further strengthen phase locking.

**Figure 6:**
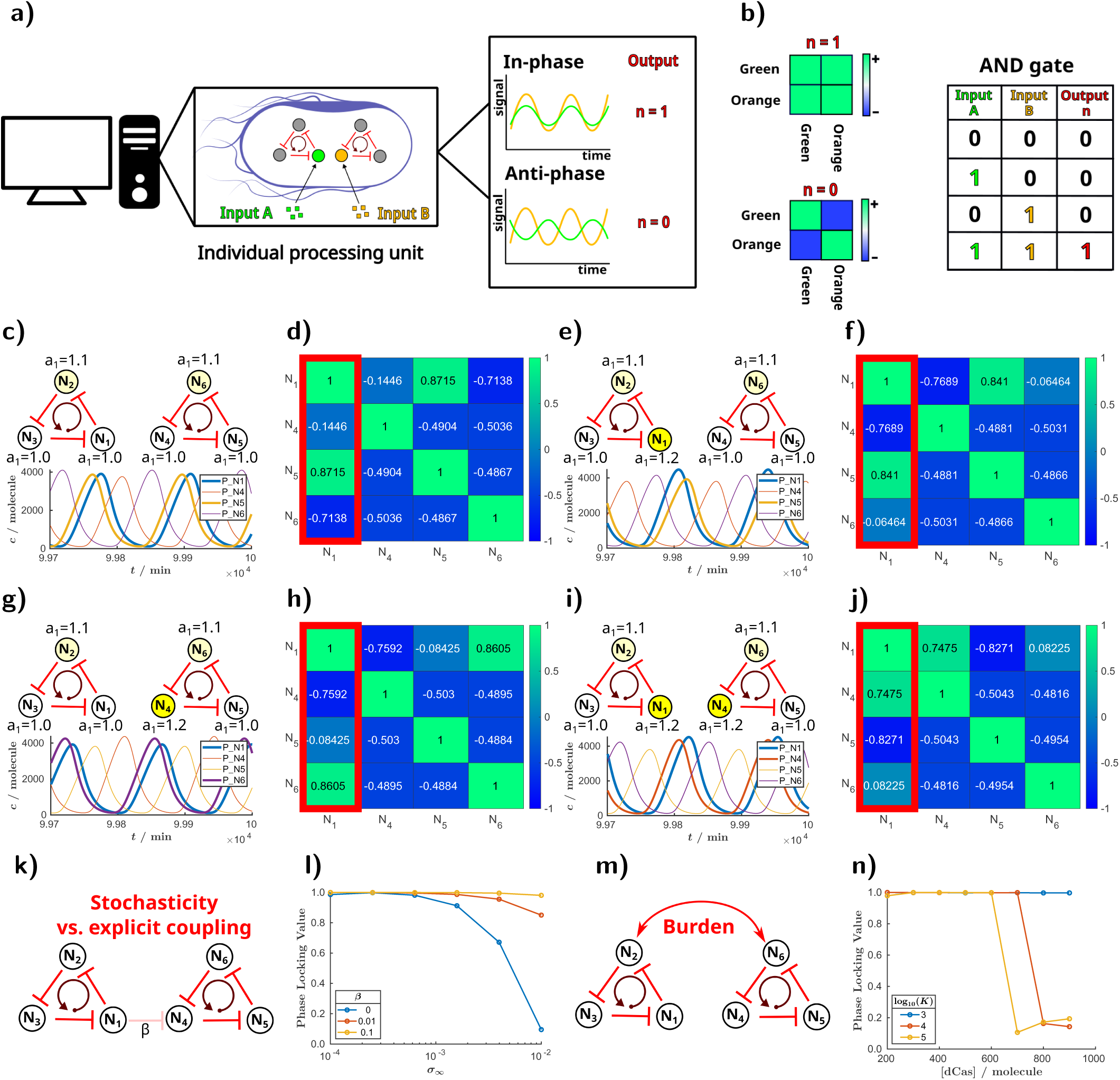
Oscillator-Based Computation: Phase Locking in Coupled CRISPRlators. **a)** Schematic of intracellularly coupled oscillators, where small chemical inputs drive binary outputs based on phase differences. **b)** Correlation matrix showing in-phase and anti-phase signals, with an example AND gate operation. **c, e, g, i)** Circuit topologies (top) and protein oscillations (bottom), demonstrating phase locking with low dCas concentration (200 molecules). **d, f, h, j)** Correlation matrices computed from protein time series. **k)** Schematic representation of an alternative (explicit) coupling strategy between oscillators. **l)** The effect of the repression strength between the oscillators (*β*) to the phase-locking. In panels (l) and (n), the simulations were run for 10^5^ minutes, with the first 15% treated as transient. Parameters were *a*_1,*N*2_ = 1., *a*_1,*N*6_ = 1.2, and in (n) [dCas] = 200 molecules. **m)** Schematic illustration of implicit coupling between oscillators mediated by the time-dependent burden. **n)** Phase-locking value versus dCas concentration for various half-saturation constants *K*; lower *K* indicates stronger burden-m_13_ediated coupling. The details of the simulations are available in “computer.html.” An extended figure caption with additional details is available in SI Section 5

During the initial oscillations, synchronization occurs, representing the phase in which the system performs its computation. The output of a computation with oscillators is not a single low or high signal intensity; instead, the information is encoded in the frequency or phase of the output. Here, we focus on the phase difference between a node in the first oscillator (N_1_) and nodes in the second oscillator (N_4_, N_5_, N_6_), while our input is the promoter strength of N_3_ and N_4_ (Figure 6). Once synchronization occurs, the phase-locking effect enables the system to function as a memory unit. Thus, the system initially operates as a processor, performing computations during the start of oscillations, and transitions into acting as a memory unit thereafter.

We first demonstrate that the system can operate in a traditional binary mode based on Boolean algebra, functioning as an *AND* gate (Figure 6a, b). The behavior depends on whether N_1_ is synchronized with N_4_. We computed the correlation matrix from the oscillatory patterns of protein concentrations at each node (Figure 6d, f, h, j and Figure S22). However, a big proportion of these matrices are redundant: by identifying the closest node in phase with N_1_, we can qualitatively infer the remaining correlations. Nodes of the same CRISPRlator always exhibit different phases and non-positive correlations. In other words, the first column of the matrix contains all the qualitative information needed to distinguish the three possible outcomes of the system. In panels (d) and (f) of Figure 6, N_1_ is synchronized with N_5_ (and consequently N_2_ with N_6_, and N_3_ with N_4_). In Figure 6h, N_1_ aligns with N_6_, and in panel (j), it locks with N_4_. A close phase relationship results in a positive value in the correlation matrix (close to one), which is interpreted as a logical one (true), while a non-positive (close to zero or negative) correlation corresponds to a logical zero (false). In this example, the inputs could be small chemical molecules or light that modulate the promoter strengths of N_1_ and N_4_. When both N_1_ and N_4_ are activated, corresponding to high concentrations of the input chemicals, they synchronize, resulting in a positive correlation. In all other cases, the correlation is non-positive, enabling the system to function as an *AND* gate, where the output is true only if both inputs are true.

We can also illustrate the behavior of these coupled oscillators in phase space on the P_N1_-P_N2_ plane (Figure S24). This representation resembles the output of a flow cytometry measurement. Interestingly, since we are plotting two periodic signals as functions of each other, we observe *Lissajous curves*. When the correlation is perfect (i.e., one) and the phase difference is zero, the result is a straight line (Figure S24h). Conversely, with a phase difference of *π*, the line has a slope of -1. For a phase difference around *π/*2 (Figure S24a), we observe a closed curve with a large enclosed area. As the phase difference approaches 0 or *π*, this curve transitions into the straight lines described earlier (Figure S24b-d). If synchronization does not occur, the output of such a flow cytometry experiment would spread across and fill the phase space (Figure S24f, g).

Interestingly, this OBC system based on two coupled CRISPRlators, is not only capable of operating in a binary mode, it can actually produce three distinct outputs dependent on node synchronization, thus behaving as a *ternary system* (Figure S21a). Specifically, the phase of node N_1_ aligns with the phase of node N_4_, N_5_, or N_6_ from the other oscillator, producing three possible outcomes. This behavior is evident in the correlation matrices of the oscillatory signals (Figure 6f, h, j), which display three distinct patterns. However, we can observe this ternary behavior by analyzing the protein oscillations of just two nodes and calculating their corresponding correlation values (Figure S21a). The correlation can take three distinct values: -1, 0, and 1; effectively forming a ternary system. The advantage of a ternary system over a binary one lies in its ability to store more information per unit. Each ternary digit represents three states, thus reducing the number of digits required to encode the same amount of information compared to binary systems. By coupling oscillators with more nodes, the number of possible outputs can be significantly increased. Furthermore, the correlation values can vary according to the promoter strength, resulting in a *nonlinear output* (Figure S21b), further enriching its dynamic behavior. The output of the system, which represents the correlation between N_1_ and N_2_, follows a sigmoid function around a promoter strength of 1.2 min*^−^*^1^. This type of function is commonly used in neural networks. If these nonlinear functions could be computed with OBC in a single step, rather than through the layered architecture of current neural networks, we could unlock a substantial performance boost in tasks like image processing, mimicking the efficiency of biological neurons (Figure S21c).

A critical consideration in the biological implementation of OBC within cells is the presence of noise, an intrinsic feature of all biological systems. To assess its influence, we conducted stochastic simulations (Section 5) while systematically varying the noise strength, *σ_∞_* and analysed the resulting phase-locking value (PLV). The PLV quantitatively captures how consistently the phase difference between two oscillators remains stable over time, with values approaching 1 indicating strong phase locking and values near 0 signifying desynchronization (Section 2, Equation S17). Our results show that increasing noise levels progressively reduce the phase-locking value (PLV) (Figure 6l, blue curve), indicating that sufficiently strong noise can disrupt oscillator coupling. Furthermore, while phase locking typically results in the phase difference fluctuating around a stable value, high noise amplitudes lead to pronounced temporal variations in phase difference (Figure S25).

Successfully implementing such circuits experimentally will require coordinated theoretical and experimental efforts to account for these effects. One strategy to mitigate the impact of noise is to increase the coupling strength between oscillators. For example, additional explicit coupling interactions can be introduced to enhance synchronization between oscillators (Figure 6k), such as repression from node N1 to N4. Ideally, the strength of these interactions could be modulated using chemical inducers, allowing fine-tuned control over oscillator coupling. In our stochastic simulations (Figure 6k,l), we implemented such repression, with the parameter *β* scaling the baseline repression strength between nodes. As *β* increases, phase locking becomes progressively more robust, persisting even under substantial noise. Remarkably, even a modest repression level (10% of the standard repression strength) is sufficient to synchronize the oscillators despite intense noise. This behavior is further illustrated by the instantaneous phase difference between oscillators (Figure S26): under high noise and weak coupling, no phase locking is observed; with moderate coupling, phase locking occurs intermittently; and with sufficiently strong coupling, the phase difference fluctuates narrowly around a constant value, indicating stable and sustained synchronization. In addition to increasing the coupling strength, the overall coherence of the system can also be improved by reducing intrinsic noise. When intracellular oscillators are coupled through diffusible signaling molecules, intercellular synchronization can emerge. In this synchronized state, the population of cells effectively acts as a single oscillator with a larger effective volume, thereby reducing stochastic fluctuations and further enhancing the relative influence of intracellular coupling [44].

Moreover, there may be couplings beyond those we design explicitly. We illustrate this effect through the burden and resource limitation imposed by the oscillators on the host cells (Figure 6m). Because gene expression and mRNA production oscillate in time, the availability of intracellular resources, such as RNA polymerases, ribosomes and overall cellular activity also vary, potentially introducing an implicit coupling between oscillators [41–43]. We examined this effect in a simple simulation (Figure 6n), assuming that increased heterologous protein concentration leads to higher burden and consequently decreases overall cellular activity, including mRNA production [41, 78, 79]. This was modeled as a simple repression using a Hill function, where the half-saturation constant *K* was varied. The results show that as the dCas concentration increases, the PLV decreases and the oscillators lose synchrony. However, reducing *K* strengthens coupling through the burden, and the oscillators remain phase-locked even at high dCas concentrations (blue curve). It is important to emphasize that this burden is only a demonstration of implicit, indirect coupling: in reality, other mechanisms could have similar effects and modify the effective coupling strength between oscillators.

In summary, with OBC we present a novel framework for biocomputing in synthetic biology that holds promise for establishing a new paradigm, though its practical realization will require addressing technical challenges.

## Discussion

Here, we computationally explored intracellularly coupled autonomous oscillators. In other words, we computationally introduced different oscillators within a single cell and aimed to map the range of behaviours that can arise in such systems. To establish a clearer framework, we categorized the oscillators into three distinct groups based on the interaction strength between them: independent, weakly/strongly coupled, and deeply coupled oscillators, with the coupling strength progressing from zero to infinity.

The novelty of this work lies in establishing a biologically grounded framework for autonomous intracellular coupling between oscillators. We thus expand the design space of synthetic circuits toward more complex yet experimentally feasible behaviors. In the following discussion, we summarize the phenomena investigated and suggest potential future applications, many of which are inspired by existing implementations of oscillators in other fields.

### Beat phenomenon

Using the example of independent oscillators, we exemplify the beat phenomenon (Figure 3I), which has numerous practical applications. In acoustics, the beat phenomenon is commonly applied to tuning musical instruments, enabling precise adjustments by comparing interference patterns [80]. Similarly, in optical frequency metrology, it plays a crucial role in comparing and calibrating laser frequencies, facilitating advancements in precision measurement technologies [81]. In a biological context, this phenomenon could be applied to measure frequency differences between oscillators, potentially leading to the creation of novel sensors. For instance, one oscillator could be sensitive to a specific chemical compound or physical parameter, while the other remains unaffected, allowing us to measure frequency shifts through the beat phenomenon. The output frequency of the system is highly sensitive to the frequency difference between the oscillators (Figure 3Id), enhancing the precision and sensitivity of the method (Figure 7a).

**Figure 7:**
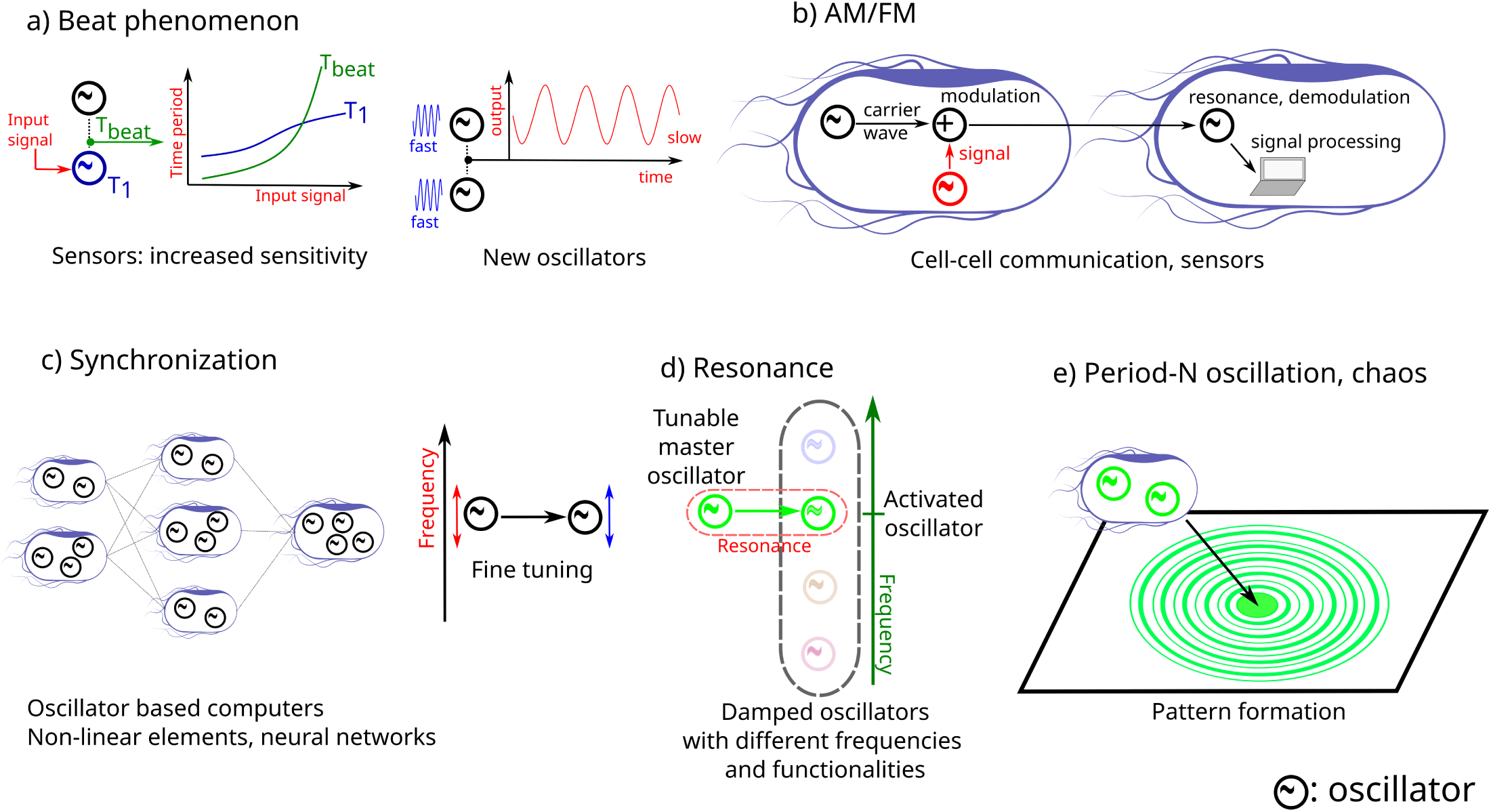
Potential applications of the phenomena demonstrated in this study. **a)** Beat phenomenon and its use in sensors and the design of new oscillators. **b)** AM and FM in cell-cell communication or as sensing mechanisms. **c)** Synchronization and oscillator-based computing, illustrated using neural network examples and a possibility to fine tune oscillators. **d)** Resonance effects for selectively activating damped oscillators, enabling new cellular functionalities. **e)** Higher-period oscillations and chaos-generated patterns. Detailed descriptions of these applications are provided in the text.

### Amplitude and frequency modulation (AM/FM)

We proposed several circuits for generating AM and FM signals. In our first example, we coupled independent oscillators with different frequencies to a common output (Figure 3II). In the second example, we used coupled oscillators (Figure S19) to generate pure AM and FM signals [56, 77]. Recent findings show that living organisms leverage this technique for cell-to-cell communication, benefiting from its enhanced signal-to-noise ratio (Figure 7b) and enabling them to accurately respond to gradual changes in external stimuli [82–85]. For example, bacteria use both AM and FM to broadcast signals that determine whether the population will commit to biofilm formation or cell motility [86]. In synthetic biology, the use of AM/FM has recently been demonstrated in mammalian cells by changing the diffusivity and reactivity of oscillatory components of a genetic circuit (based on bacterial MinDE), that can ultimately modulate the frequency and amplitude of oscillations within cells [87].

AM is commonly used in signal transmission, such as in telecommunications [5], various types of advanced sensors, including Quartz Crystal Microbalance (QCM) sensors [88], atomic force microscopy [89], electroencephalogram (EEG) [90], dielectric characterization of solids and microfluidics [91], and biomedical sensors in cochlear implants and in cardiac pacemakers [7]. AM enhances these sensors’ ability to detect subtle changes, increasing sensitivity and accuracy. Inspired by this, a potential application of AM in SynBio would be its use in whole cell biosensors [92], for applications as diverse as detecting polychlorinated biphenyls in wastewater, metallic trace elements in contaminated soil, or hallmark molecules of infectious diseases in the medical field.

### Synchronization and Oscillator-Based Computing

Synchronization was first observed and described by Huygens in the 17th century, using the example of pendulum clocks that naturally synchronized when placed on the same surface [11]. In our simulations, we reproduced that oscillators with the same frequency can shift their phase relative to each other, while oscillators with different frequencies can adjust their time periods until they synchronize depending on the strength of the coupling. In a cell, oscillators are never completely independent, which opens up the possibility of fine-tuning natural oscillators without directly modifying them. By introducing a synthetic oscillator with a frequency similar to that of a natural oscillator, we could potentially adjust the frequency and phase of a natural oscillator (cell cycle, circadian clock, metabolic oscillation) through synchronization with a synthetic oscillator.

In our numerical example of OBCs, we demonstrated how coupled CRISPRlators can be used to construct logic gates and memory units. However, the potential of OBCs extends far beyond these applications. By moving beyond binary thresholds and Boolean logic and instead utilizing partial synchronization between oscillators, it becomes possible to generate nonlinear functions as output. This approach could prove valuable in tasks such as neural network computations, image analysis (Figure S21c) [66, 93, 94] or in oscillator-based optimization [95]. Moreover, coupling more than just two oscillators significantly increases the number of possible states, potentially enabling solutions to computationally hard problems in a manner analogous to entangled systems in quantum information theory [8, 66]. However, scaling up the number of coupled oscillators will require intercellular coupling and the development of stable co-cultures [39, 96–98], which represents an important next step in this field. In our example (Figure 6), we observed that information can be stored in the phase of oscillators through phase locking. Alternatively, the information could also be encoded in the frequency of the oscillator [66]. Nature already employs this principle of frequency encoding. A striking example is found in snakes, whose pit organs can detect temperature differences as small as a few millikelvins [99]. In this system, sensory information is carried by a stochastic oscillator: minute temperature shifts, particularly when the system operates near a bifurcation point, can induce large changes in firing frequency. This phenomenon illustrates how stochasticity does not merely disrupt OBC, but can instead be exploited to achieve exceptional sensitivity. By tuning the system close to a critical transition, weak inputs can be amplified into robust frequency shifts. Building on this idea, we next discuss how resonance and coupling between oscillators could be leveraged to read out frequency-encoded signals in synthetic systems.

### Resonance

Previous studies demonstrated that in a non-autonomous system, when the frequency of external forcing matches the natural frequency of the synthetic oscillator, high-amplitude oscillations, i.e., resonance, can be observed [33, 34]. In this work, we show that a unidirectional coupling between autonomous oscillators produces an analogous behavior, resulting in resonance as well (Figure 5).

Resonance is widely utilized in electrical engineering, such as in radio receivers or band-pass filters, where only a specific frequency range is targeted. In certain methods, resonance is advantageous, such as in Nuclear Magnetic Resonance (NMR) or Electron Paramagnetic Resonance (EPR) [100]. However, in structures such as tall buildings, resonance-induced vibrations are mitigated by using tuned mass dampers [101]. In SynBio, oscillatory circuits often exhibit damped oscillatory behavior rather than sustained oscillations. As a potential application, introducing another robust oscillator into the system could drive the damped oscillator at its resonant frequency, forcing it to oscillate. By incorporating multiple damped oscillators with varying natural frequencies and properties, we could harness their resonance and band-pass filter characteristics to selectively activate specific oscillators by tuning the frequency of a master oscillator. This approach ensures that only the activated damped oscillator oscillates, reducing the cellular burden while enabling the desired functionality associated with the selected oscillator (Figure 7d). Furthermore, the system can be easily fine-tuned by adjusting the master oscillator’s frequency with a single inducer. In another application, resonance could be harnessed to filter and amplify AM and FM signals in cell-to-cell communication.

### Period-*N* oscillation, chaos

In SynBio, oscillators have been used to generate spatial patterns [28–30, 33]. Bacterial colonies harboring a synthetic oscillator and growing on solid surfaces produce ring patterns. These patterns are formed because only the cells at the colony edge are growing and oscillating, while cells in the inner region are arrested in different phases of the oscillations. We previously demonstrated that using higher-period oscillations, we can produce rings with alternating intensities, and that these patterns are disrupted in chaotic regions [33]. However, in that case, an externally modulated light signal was required to achieve the effect. With the suggested autonomous chaotic coupled oscillators, it could be possible to generate these patterns without the need for an external signal (Figure 7e).

Surprisingly, we found that the most well-known repressilator and toggle switch circuits should be able to exhibit Goodwin-type oscillations, caused by enzymatic protein degradation (Figure S13), SI Section 3.2). In this oscillatory state, nodes synchronize, leading to fast, low-amplitude oscillations. Due to the interactions between different oscillatory states, we were even able to predict chaotic behavior in a small parameter region. A noteworthy experimental implication is that if these fast, low-amplitude oscillations were observed without prior knowledge of this phenomenon, it might be mistaken for malfunctioning behavior, prompting unnecessary circuit modifications. This underscores the importance of theoretical mapping to complement experiments, as even well-characterized circuits can exhibit unexpected behaviors. Learning these possibilities beforehand allows for a better understanding of experimental outcomes and avoids misinterpretation.

### Future work

The main objective of this study was to explore the potential of intracellularly coupled oscillators. While this work serves as a foundation, we hope it will encourage further computational and experimental studies. For example, studying intracellularly coupled oscillators can deepen our understanding of the mechanisms and interactions that govern ubiquitous natural oscillators. In addition, this research may yield critical insights into how synthetic oscillators interact with their natural counterparts. Such understanding could provide a powerful tool for engineering living systems, enabling the manipulation of biological processes through the coupling of synthetic and natural oscillators.

Further research is also required to experimentally implement synthetic intracellularly coupled oscillators. For instance, for deeply coupled oscillators, we showed that the same phenomena can be achieved through various methods, such as altering promoter strengths, protease concentrations, or repression strengths. Experimental realization will require identifying the optimal method to construct these circuits, as this study focused solely on fundamental phenomena.

However, we are optimistic that implementing coupled oscillators in *E. coli* is feasible. Prindle and colleagues, for instance, have already provided a proof of concept by successfully coupling two oscillators to achieve frequency modulation [44]. Regarding noise, it is worth emphasizing that individual oscillators serving as building blocks have been highly optimized to exhibit remarkably robust dynamics with minimal noise and phase drift [30–32, 102, 103]. For example, certain repressilator variants require up to 455 generations to drift by half a phase [102]. Furthermore, oscillator frequency can be externally modulated using small molecules or light [33–35], enabling systematic parameter exploration and fine-tuning. Multiple orthogonal oscillators without shared parts have also been successfully implemented [102].

A potential concern is the metabolic burden imposed by multiple oscillators on the host cell. Yet, a five-node re-pressilator has already been demonstrated, and the addition of one or two nodes represents only a modest increase in network size. More strikingly, a synthetic circuit comprising 35 genes (31 kb of DNA) has recently been implemented in a single *E. coli* cell, suggesting that accommodating two relatively small oscillators is well within reach. Moreover, CRISPRi-based circuits offer a promising alternative to transcription factor–based oscillators, as orthogonal sgRNAs can be readily designed, and their transcription generally imposes a low cellular burden [104]. We have previously shown that *E. coli* can host two functional three-node networks simultaneously [31]. While high levels of dCas9 can be toxic [30–32, 102, 103], this limitation can be mitigated using engineered dCas9 variants or by employing dCas12a, which has been reported to be less toxic [105, 106].

A central challenge in experiments with coupled oscillators will be the need to record a sufficient number of oscillatory cycles to reliably characterize system dynamics. Since most synthetic oscillators depend on actively growing cells, experimental setups that allow to maintain long-term cultures will be required. Employing faster oscillators, such as the Goodwin oscillator, could be advantageous in this context. At these extended timescales, optimising the evolutionary stability of the circuits may also become necessary.

When considering real-world applications of those circuits, environmental factors, notably temperature, must also be taken into account. The frequency of synthetic oscillators is known to be temperature-sensitive, which can affect their performance and dynamic behavior [26, 107]. One way to address this limitation is through the design of temperature-compensated circuits, in which oscillatory periods remain stable despite fluctuations in temperature. The circadian clock provides a natural example of such robustness [108–110], and Hussain and colleagues have shown that temperature compensation is also achievable in a synthetic oscillator [111].

Another critical factor in these systems is the coupling strength. For independent or deeply coupled oscillators, determining the coupling mechanism is relatively straightforward, but for weakly or strongly coupled oscillators, careful consideration of coupling strength is essential. We proposed two potential mechanisms: protease-mediated coupling (Figure 5) and coupling via the dCas system (Figure 6). However, coupling could also emerge from other limited cellular resources [41–50] (Figure 6). In cases where the origin of synchronization is unclear, the well-established Kuramoto model [39, 112, 113] could serve as a useful framework for studying coupled oscillators. Finally, in OBC systems, computation occurs during the initial phase of oscillation, yet our understanding of this transient phase remains limited. This gap highlights the need for additional experiments and possibly the development of new models.

Biological noise also modulates the coupling strength necessary for stable synchronization. As shown by our stochastic simulations, higher noise levels lead to a progressive decline in phase-locking (Figure 6l), indicating that strong noise can disrupt coupling. However, coupling can be restored either by incorporating additional explicit interactions or through coupling mediated by limited shared resources (Figure 6k-n). Thus, our findings suggest that OBC remains feasible and robust even under biologically noisy conditions.

The proposed systems operate on timescales of minutes to hours, which raises the question of whether such dynamics are sufficiently fast for real-time applications. However, several existing synthetic biology platforms already perform computation on comparable or even slower timescales [58, 114], and yet interest in this field is expanding rapidly because of its transformative potential. OBCs may help overcome this limitation by leveraging coupling and synchronization to enable inherently parallel computation, thereby offering the prospect of significant acceleration. Theoretical analyses [66] also indicate that OBCs could solve computationally complex problems more efficiently than conventional digital systems. In such cases, slower timescales may be an acceptable trade-off for enhanced computational capability. Although these prospects remain speculative, we anticipate that ongoing conceptual and technological advances will continue to stimulate progress in this emerging area.

In conclusion, we believe that intracellularly coupled oscillators hold considerable promise for synthetic biology, both as a fundamental design principle and as a platform for diverse applications, making them a compelling direction for future research.

## Methods

We used the GRN modeler [55] to design and model the circuits. Simulations were conducted in MATLAB (version 2023b) using the SimBiology toolbox with the Sundials solver. The simulations were performed with relative tolerances set to 10*^−^*^8^ and absolute tolerances set to 10*^−^*^10^, unless stated otherwise in the figure captions.

We determined the time period (*T* ) of the oscillators and, for the beat phenomenon in Figure 3, calculated the standard deviation of *T* using MATLAB’s findpeaks function.

### Bifurcation diagrams

#### Linear stability analyzis

During the linear stability analysis (SI, Section 3.2) of the Goodwin oscillator, the fixed point was determined using MATLAB’s `fsolve` function, where the right-hand side of the ODE system was set to zero. The Jacobian matrix was then evaluated at the fixed point. If the real part of all eigenvalues of the Jacobian was negative, the fixed point was considered stable. For cases where the fixed point was unstable and had complex eigenvalues, stable periodic orbits were identified by starting simulations from the unstable fixed point. A 2 *·* 10^3^ minute simulation was run, and the highest and lowest values of the protein oscillations were calculated from the second half of the simulation.

In the case of the toggle switch a similar procedure was performed; however, we had to check the stability of multiple fixed points. The fixed points were found first analytically with the `solve` function, then it was evaluated with the `vpa` function at the given protease concentrations. The stability of the fixed point was determined based on the eigenvalues of the Jacobian matrix.

#### Calculating the Lyapunov exponents

In the bifurcation diagrams for the visualization of the SPOs and the chaotic regions, each point represents the outcome of a simulation lasting 2 *·* 10^5^ minutes. The first half of the simulation was discarded as transient behavior, while the second half was used to construct the bifurcation diagram. Following this, the simulation was extended to compute the Lyapunov exponents. These exponents were calculated using COPASI [115] via the BasiCo interface [116] in MATLAB. Specifically, after a preliminary 10^3^ minute simulation, the Lyapunov exponents were determined from a subsequent 10^4^ minute simulation, with orthogonalization performed every 10^2^ minutes. For these calculations, a relative tolerance of 10*^−^*^4^ and an absolute tolerance of 10*^−^*^6^ were applied. To accurately identify positive Lyapunov exponents and differentiate them from near-zero values, we computed the standard deviation (*σ*) of the exponents closest to zero. Exponents were considered positive if they exceeded 3*σ*, ensuring that the regions in the bifurcation diagrams marked with red (indicating positive Lyapunov exponents) were not artifacts of small numerical errors, particularly when theoretical values should be zero.

## Code availability

This code is based on the *GRN modeler* platform [55], which provides tools for constructing, simulating, and analyzing gene regulatory networks. The code used to generate the results presented in this study is available at the following GitHub repository: https://github.com/SchaerliLab/GRN_modeler/tree/main/analysing_examples/coupled_oscillators.

## Acknowledgments

This work was funded by the Swiss National Science Foundation (200532 and 10005200 awarded to Y.S.), a fellowship of the Agassiz foundation (awarded to J.P.), a UNIL FBM Ph.D fellowship in Life Sciences (awarded to J.P.) and the University of Lausanne.

## Author contributions

G.H. performed the mathematical modeling. G.H., J.P., R.E., and Y.S. wrote the manuscript. All authors have given approval to the final version of the manuscript.

## Supplementary information

Supplementary Information.pdf, AM.html, deeply coupled Tomazou.html, AM independent.html, FM AM.html, AM independent Stricker weak.html, FM.html, beat.html, goodwin.html, computer.html, repressilator goodwin.html, coupled repressilators Tomazou.html, repressilator repressilator.html, deeply coupled Elowitz.html, repressilator Tomazou.html, deeply coupled Elowitz inducer.html, resonance.html.

