## Supplementary Information for "Intracellularly Coupled Oscillators for Synthetic Biology"

### 1 The applied models

In this section, we summarize the foundations of the models applied in our study. All simulations were performed using the *GRN_modeler* application [1], which incorporates these models. For the reader's convenience, the following tables and accompanying text have been adapted from our previous publication [1]. They present key parameters and model definitions essential for understanding the simulations. We include them here in the Supplementary Information with identical content to ensure completeness, clarity, and ease of reference.

#### 1.1 Elowitz-model

The specifics of the Elowitz model are provided in Table S1. In reaction $R_1$, the $i$th node generates mRNA, which can be inhibited by a transcription factor originating from the $j$th node. Reaction $R_2$ describes the production of the transcription factor associated with the $i$th node.

**Table S1:** The Elowitz-type transcription factor model [1, 2] describes the production of mRNA and protein ($mRNA_i$, $P_i$) at the $i$th node, which is repressed by another transcription factor, $P_j$. This repression is modeled using a Hill function: $HILL([P_j]) = \frac{1}{1 + ([P_j]/K)^n}$, where K = 40 molecule and n = 2.

| Nr. | Reaction | Rate law | Rate constant [2] | Unit |
|---|---|---|---|---|
| $R_1$ | $\emptyset \longleftrightarrow mRNA_i$ | $r_1 = k_0 + k_1 \cdot HILL([P_j])$ | $k_0 = 0.03$ | molecule/minute |
| | | | $k_1 = 30$ | molecule/minute |
| | | $r_{1r} = k_2\,[mRNA_i]$ | $k_2 = 0.3466$ | 1/ minute |
| $R_2$ | $\emptyset \longleftrightarrow P_i$ | $r_2 = k_3\,[mRNA_i]$ | $k_3 = 6.9315$ | 1/ minute |
| | | $r_{2r} = k_4\,[P_i]$ | $k_4 = 0.0693$ | 1/ minute |

#### 1.2 Tomazou-model

In Table S2 we present the Tomazou-type transcription factor model [3]. The mRNA production of the $i$th node can be repressed by the transcription factor of the $j$th node, $P_j$. This repression is described with the following Hill-function: $HILL([P_j]) = \frac{1}{1 + ([P_j]/K)^n}$, where K = 5 molecule and n = 2. When external inducers are present ($I_1$ and $I_2$, $[P_j]$ can be replaced with the concentration of the inducers in the Hill-function and K = 50 $\mu$M. The protease degradation rate is determined using Michaelis-Menten kinetics: $k_{protease} = \frac{k_{protease,max}[PROT]}{K_{protease} + Substrates}$, where $k_{protease,max} = 50$ 1/ minute, $K_{protease} = 30$ molecule and "Substrates" represents the sum of all proteins degraded by the given protease ($Substrates = \sum_i [P_i]$).

**Table S2:** The Tomazou-type transcription factor model [1, 3]. Further explanation can be found in the text.

| Nr. | Reaction | Rate law | Rate constant [3] | Unit |
|---|---|---|---|---|
| $R_1$ | $\emptyset \longleftrightarrow mRNA_i$ | $r_1 = n_{copy}(a_0 + a_1 \cdot HILL([P_j]))$ | $n_{copy} = 25$ | molecule |
| | | | $a_0 = 0.001$ | 1/ minute |
| | | | $a_1 = 100$ | 1/ minute |
| | | $r_{1r} = (k_{mRNA,degr} + k_d)\,[mRNA_i]$ | $k_{mRNA,degr} = 0.5$ | 1/ minute |
| $R_2$ | $\emptyset \longleftrightarrow uP_i$ | $r_2 = k_{translation}\,[mRNA_i]$ | $k_{translation} = 6.9315$ | 1/ minute |
| | | $r_{2r} = (k_d + k_{protease,i})\,[uP_i]$ | $k_d = 0.01$ | 1/minute |
| $R_3$ | $uP_i \longrightarrow P_i$ | $r_3 = k_{mat}\,[uP_i]$ | $k_{mat} = 0.4$ | 1/ minute |
| $R_4$ | $P_i \longrightarrow \emptyset$ | $r_4 = (k_d + k_{protease,i})\,[P_i]$ | | |

##### 1.3 CRISPR model

**Table S3:** Modified model for the CRISPRlator. Initial concentrations are set at $[DNA]_0 = 30$ molecules and $[dCas]_0 = 1434$ molecules [1, 4]. All rate constants are provided in units of *minute* and *molecule*. The repression of the $j$th node on the $i$th node occurs through the formation of the $DNA_j$-$sgRNA_i$ complex, as described in the $R_6$ reaction.

| Nr. | Reaction | Rate law | Rate constant | Ref. |
|---|---|---|---|---|
| $R_1$ | $\emptyset \longleftrightarrow mRNA_i$ | $r_1 = a_0 + a_1\,[DNA_i]$ | $a_0 = 0.03$ | [2] |
| | | $r_{1r} = (k_d + d_{RNA})[mRNA_i]$ | $a_1 = 1$ | |
| $R_2$ | $\emptyset \longleftrightarrow sgRNA_i$ | $r_2 = a_0 + a_1\,[DNA_i]$ | $d_{RNA} = 0.3286$ | [2] |
| | | $r_{2r} = (k_d + d_{RNA})[sgRNA_i]$ | $k_d = 0.018$ | |
| $R_3$ | $\emptyset \longleftrightarrow P_i$ | $r_3 = k_P\,[mRNA_i]$ | $k_P = 6.9315$ | [2] |
| | | $r_{3r} = (k_d + d_P)[P_i]$ | $d_P = 0.0513$ | |
| $R_4$ | $dCas + sgRNA_i \longleftrightarrow dCas{:}sgRNA_i$ | $r_4 = k_{fds}[dCas][sgRNA_i]$ | $k_{fds} = 1.4674$ | [4] |
| | | $r_{4r} = k_{rds}[dCas{:}sgRNA_i]$ | $k_{rds} = 0.0776$ | |
| $R_5$ | $dCas{:}sgRNA_i \longrightarrow dCas$ | $r_5 = k_d[dCas{:}sgRNA_i]$ | | [4] |
| $R_6$ | $dCas{:}sgRNA_i + DNA_j \longleftrightarrow dCas{:}sgRNA_i{:}DNA_j$ | $r_6 = k_{fdsd}[dCas{:}sgRNA_i][DNA_j]$ | $k_{fdsd} = 0.2670$ | [4] |
| | | $r_{6r} = k_{rdsd}[dCas{:}sgRNA_i{:}DNA_j]$ | $k_{rdsd} = 0$ | |
| $R_7$ | $dCas{:}sgRNA_i{:}DNA_j \longrightarrow dCas + DNA_j$ | $r_7 = k_d[dCas{:}sgRNA_i{:}DNA_j]$ | | [4] |

#### 2 Independent oscillators

##### 2.1 Foundations of Oscillatory Signal Modulation

To aid in interpreting the features observed in the main text, we summarize analytic results for two related effects. For the beat phenomenon, we show how the time-domain superposition of two nearby-frequency harmonic oscillations produces a slowly varying envelope and a beat frequency. For amplitude modulation, we derive the Fourier spectrum of a carrier whose amplitude is modulated by a slower oscillation, yielding a central peak with symmetric sidebands. In addition, we illustrate these effects using Hilbert and wavelet transforms on these simple examples, which provide an intuitive time–frequency view and help to understand the origin of the low-frequency envelope and the spectral structure observed in the simulations.

##### 2.1.1 Beat phenomenon with simple harmonic functions

To illustrate the phenomenon observed in independently coupled oscillators, one can approximate their output signals using simple harmonic functions:

$$s_1(t) = A_1 \cos(\omega_1 t), \tag{S1}$$

$$s_2(t) = A_2 \cos(\omega_2 t), \tag{S2}$$

where $s_1(t)$ and $s_2(t)$ are the outputs of the two oscillators, modeled as cosine functions with amplitudes $A_1$, $A_2$, and angular frequencies $\omega_1$, $\omega_2$, respectively. Assuming unit amplitudes for simplicity, the sum of the two signals becomes:

$$s_1(t) + s_2(t) = 2\cos\left(\frac{\omega_1 - \omega_2}{2} t\right) \cos\left(\frac{\omega_1 + \omega_2}{2} t\right). \tag{S3}$$

This expression reveals that the resulting signal oscillates at an angular frequency equal to the average $(\omega_1 + \omega_2)/2$, while its amplitude is modulated by an envelope oscillating at the *beat angular frequency*, given by:

$$\omega_{\text{beat}} = \left| \frac{\omega_1 - \omega_2}{2} \right|. \tag{S4}$$

This slow modulation in amplitude is characteristic of beat phenomena and is closely related to the amplitude modulation observed in more complex coupled oscillator systems (see also Equation 3, Figure 3). These functions are illustrated in Figure S1.

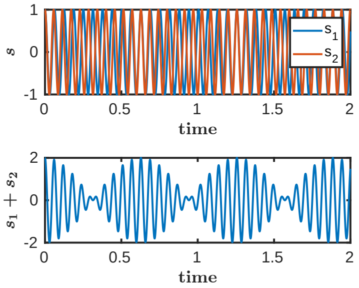

**Figure S1: Beat phenomenon.** The signals correspond to Equations S1 and S2, with $A_1 = A_2 = 1$, $\omega_1 = 100$, and $\omega_2 = 110$. The envelope of the summed signal varies at a lower frequency than the signal itself; this slow modulation is known as the *beat phenomenon*.

##### 2.1.2 Hilbert transform for signal analysis

The *Hilbert transform* is defined as the Cauchy principal value (p.v.) of the convolution of a real-valued signal with the function $\frac{1}{\pi t}$, which induces a 90° phase shift in each frequency component of the signal. This transformation is widely used to extract properties from oscillatory or modulated signals, such as amplitude, phase, instantaneous frequency, and phase difference. We illustrate its usefulness on the beat phenomenon generated by the sum of two harmonic oscillations (see Figure S2).

The Hilbert transform of a signal $s(t)$ is given by:

$$\hat{s}(t) = \mathcal{H}[s(t)] = \frac{1}{\pi}\,\text{p.v.}\int_{-\infty}^{\infty} \frac{s(\tau)}{t-\tau}\,\mathrm{d}\tau, \tag{S5}$$

where $\hat{s}(t)$ denotes the Hilbert transform of $s(t)$, and $\mathcal{H}[\cdot]$ represents the Hilbert operator.

Using this, we construct a complex-valued signal as:

$$z(t) = s(t) + i\,\hat{s}(t), \tag{S6}$$

which enables the computation of several useful quantities. The instantaneous amplitude of the signal is given by

$$A(t) = |z(t)|, \tag{S7}$$

while the instantaneous phase is defined as

$$\phi(t) = \arg(z(t)) = \tan^{-1}\frac{\mathcal{H}[s(t)]}{s(t)}. \tag{S8}$$

From the phase, we can compute the instantaneous frequency:

$$\nu(t) = \frac{1}{2\pi}\frac{\mathrm{d}\phi(t)}{\mathrm{d}t}. \tag{S9}$$

Furthermore, for two signals with respective phases $\phi_1(t)$ and $\phi_2(t)$, the instantaneous phase difference is:

$$\Delta\phi(t) = \phi_1(t) - \phi_2(t). \tag{S10}$$

This phase difference serves as a key indicator of synchronization: it is constant if the oscillators are phase-locked, and time-varying if they are not.

As an example, consider the signal formed by the sum of two cosine functions with different frequencies ((Equation S3). The analytical amplitude of the beat signal is:

$$A(t) = 2\left|\cos\left(\frac{\omega_1-\omega_2}{2}t\right)\right|. \tag{S11}$$

The approximate phase of the signal is:

$$\phi(t) = \frac{\omega_1+\omega_2}{2}t, \tag{S12}$$

which leads to a constant instantaneous frequency:

$$\nu = \frac{\omega_1+\omega_2}{4\pi}. \tag{S13}$$

The instantaneous phase difference between the two original signals is:

$$\Delta\phi(t) = (\omega_1-\omega_2)\,t, \tag{S14}$$

which increases linearly over time, indicating the absence of synchronization. This example illustrates how the Hilbert transform can be used in numerical simulations and even in experimental analyses (Figure S2).

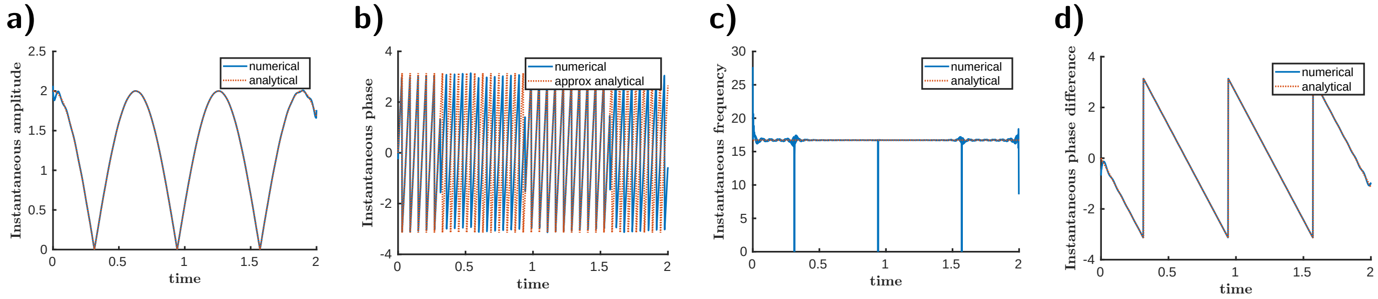

**Figure S2: Illustrating the extraction of signal properties using the Hilbert transform.** The time interval $[0, 2]$ was divided into 1000 equidistant points. We compared the numerically extracted instantaneous **(a)** amplitude, **(b)** phase, and **(c)** frequency of the mixed signal (Equation S3), as well as the **(d)** phase difference between the two oscillators, with the corresponding analytical expressions.

###### 2.1.3 Wavelet transform for signal analysis

The Hilbert transform is commonly used to extract instantaneous phase and amplitude from oscillatory signals. However, it becomes unreliable when the signal contains plateaus (what we can observe sometimes in our simulations). In such cases, the continuous wavelet transform (CWT) provides a robust alternative. The CWT of a signal $x(t)$ is defined as:

$$W_x(a,b) = \frac{1}{\sqrt{a}} \int_{-\infty}^{\infty} x(t)\, \psi^* \left( \frac{t-b}{a} \right) \mathrm{d}t, \tag{S15}$$

where $\psi(t)$ is the mother wavelet, $a$ is the scale parameter (related to frequency), and $b$ is the time shift. The complex-valued coefficients $W_x(a,b)$ allow extraction of both instantaneous amplitude and phase at each time-frequency point.

By comparing the CWTs of two input signals, one can visually assess synchronization by examining the alignment of dominant frequencies over time. To quantify phase synchronization, we calculate the *instantaneous phase difference*:

$$\Delta\phi(t) = \phi_1(t) - \phi_2(t), \tag{S16}$$

where $\phi_1(t)$ and $\phi_2(t)$ are the phases obtained from the wavelet transform of each signal. Then we calculate the *phase-locking value (PLV)*, which provides a quantitative measure of phase synchronization:

$$\mathrm{PLV} = \left| \frac{1}{N} \sum_{n=1}^{N} e^{i\Delta\phi(t_n)} \right|, \tag{S17}$$

where $N$ is the number of time points. A PLV near 1 indicates strong phase locking, while values near 0 indicate phase independence.

We illustrate CWT on our previous example analyzing the beat phenomenon (Figure 3). Figures S3a and b show that the input signals maintain constant frequency over time, indicating the absence of frequency synchronization. In contrast, panel (c) reveals the emergence of beats in the output signal. Panel (d) displays the PLV, which remains low across all frequencies, confirming that the signals are independent and that no phase locking occurs in the system.

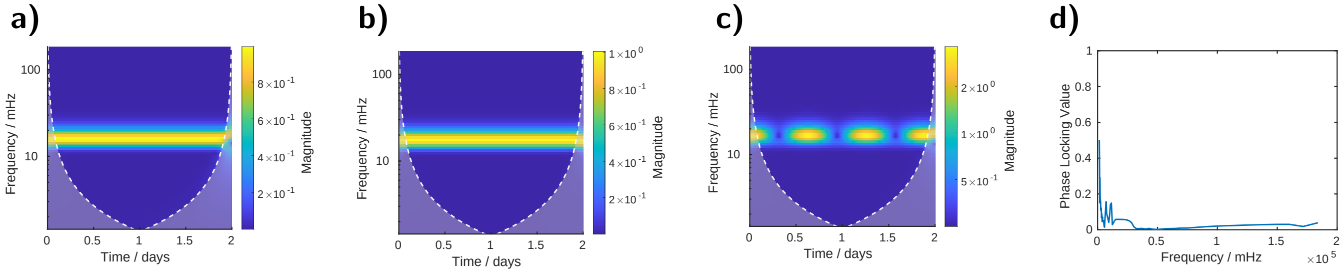

**Figure S3: Illustrating the extraction of signal properties using the Wavelet transform. a–c)** Continuous wavelet transforms of the signals defined by Equations S1, S2, and S3, respectively. **d)** Phase-locking value as a function of frequency, computed from the phase difference between the CWT representations shown in panels (a) and (b).

###### 2.1.4 Hilbert transform for non sinusoidal signals

In practical applications, oscillatory signals are often non-sinusoidal, which can complicate the use of the Hilbert transform. To extract the instantaneous phase from such signals, which may exhibit sharp rises or rapid transitions, we first estimated the average oscillation period $T$ using zero-crossing analysis. Then the signal was filtered around its fundamental frequency $f_0 = 1/T$ using a narrow bandpass Butterworth filter. The normalized cutoff frequencies are

$$\omega_{\text{low}} = \frac{\max(0, f_0 - \text{BW}/2)}{f_s/2}, \quad \omega_{\text{high}} = \frac{\min(f_s/2, f_0 + \text{BW}/2)}{f_s/2}, \tag{S18}$$

where $f_s$ is the sampling rate and $\text{BW} = 0.5 f_0$ is the bandwidth.

The digital 3rd-order Butterworth filter was created by the `butter` function of MATLAB. The filtered signal is then obtained by zero-phase forward–backward filtering using the `filtfilt` function of MATLAB. This procedure preserves the phase of the oscillatory component, enabling accurate extraction of the instantaneous phase via the Hilbert transform.

###### 2.1.5 Amplitude modulation with simple harmonic functions

A straightforward way to study the fundamental properties of amplitude modulation (AM) is to vary the amplitude of a harmonic (carrier) signal with another harmonic (modulating) signal. This can be expressed as:

$$s(t) = [1 + m\cos(\omega_m t)]\cos(\omega_c t), \tag{S19}$$

where $\omega_c$ is the angular frequency of the carrier signal, $\omega_m$ is the angular frequency of the modulating signal, and $m \in [0, 1]$ is the modulation index.

Expanding this product using trigonometric identities gives:

$$s(t) = \cos(\omega_c t) + m\cos(\omega_m t)\cos(\omega_c t) \tag{S20}$$

$$= \cos(\omega_c t) + \frac{m}{2}\left[\cos((\omega_c + \omega_m)t) + \cos((\omega_c - \omega_m)t)\right]. \tag{S21}$$

To analyze the frequency components present in the system, we apply the Fourier transform to the signal. The continuous-time Fourier transform $S(\omega)$ of a signal $s(t)$ is defined as:

$$S(\omega) = \mathcal{F}[s(t)] = \int_{-\infty}^{\infty} s(t)\, e^{-i\omega t}\, \mathrm{d}t. \tag{S22}$$

We compute the transform of each cosine term individually. Using Euler's identity:

$$\cos(\omega t) = \frac{1}{2}\left(e^{i\omega t} + e^{-i\omega t}\right),$$

we find:

$$\mathcal{F}[\cos(\omega_0 t)] = \frac{1}{2}\left[\int_{-\infty}^{\infty} e^{i(\omega_0-\omega)t}\,dt + \int_{-\infty}^{\infty} e^{-i(\omega_0+\omega)t}\,\mathrm{d}t\right]. \tag{S23}$$

Applying the identity

$$\int_{-\infty}^{\infty} e^{i\alpha t}\,\mathrm{d}t = 2\pi\delta(\alpha),$$

we obtain:

$$\mathcal{F}\left[\cos(\omega_0 t)\right] = \pi\left[\delta(\omega-\omega_0) + \delta(\omega+\omega_0)\right]. \tag{S24}$$

Therefore, the Fourier transform of the AM signal in Equation S19 becomes:

$$\begin{aligned} S(\omega) = \pi\big[&\delta(\omega-\omega_c) + \delta(\omega+\omega_c) \\ &+ \frac{m}{2}\left(\delta(\omega-\omega_c-\omega_m) + \delta(\omega+\omega_c+\omega_m)\right) \\ &+ \frac{m}{2}\left(\delta(\omega-\omega_c+\omega_m) + \delta(\omega+\omega_c-\omega_m)\right)\big]. \end{aligned} \tag{S25}$$

This result confirms that amplitude modulation produces a frequency spectrum composed of the carrier frequency $\omega_c$ and two symmetric sidebands at $\omega_c \pm \omega_m$, which is a defining feature of AM signals. Although the time-domain signal displays a slowly varying envelope (i.e., the beat-like behavior), its frequency-domain structure contains only discrete spectral lines (Figure S4.

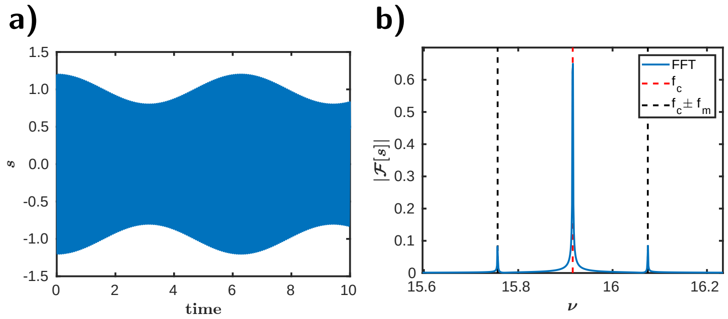

**Figure S4: Amplitude modulation.** The signal corresponds to Equation S19, with $\omega_m = 1$, $\omega_c = 100$, and modulation index $m = 0.2$. **a)** Time-domain signal. **b)** Frequency spectrum, showing a peak at the carrier frequency $\omega_c$ and sidebands at $\omega_c \pm \omega_m$, characteristic of amplitude modulation. In this simple example, the analytically and numerically computed spectra coincide perfectly.

**Extended caption for Figure 3**: **Independently coupled oscillators.**

**Row I: The Beat Phenomenon. I,a)** The topology of the circuits showing two repressilators connected via a NOR gate at node $N_7$. $PROT_1$ and $PROT_2$ represent proteases. We varied $PROT_1$ levels from 10 to 200 molecules, while keeping $PROT_2$ constant at 100 molecules. **I,b)** Trajectories of protein concentrations for the first (blue curve) and fourth (red curve) nodes, when $PROT_1$ was set to 164 molecules. The first part of the graph represents the scenario where the peaks do not overlap, resulting in continuous repression of $N_7$. In the second half, however, the peaks overlap, leading to the activation of $N_7$. **I,c)** Oscillations in the protein concentration of the output node ($N_7$). Peaks are observed when the oscillations in (b) are in-phase (second half of the graph). Conversely, when the oscillations in (b) are out-of-phase (first half), $N_7$ remains repressed. Gray rectangles indicate the time range corresponding to panel b. **I,d)** Theoretical output of the beat phenomenon (blue curve) and the simulated time period of the output node (red curve) as a function of the first repressilator's time period, with the time period of the second oscillator kept constant at 152.5 minutes (gray vertical dashed line). The simulations lasted for $10^4$ minutes, with the first $10^3$ minutes considered as transient time. The theoretical and simulated curves show strong agreement, with the simulated curve closely aligning and overlapping the theoretical one.

**Row II: Amplitude Modulation (AM). II,a)** The topology of the circuits, with a CRISPRlator and a dual-feedback oscillator connected via a NOR gate at node $N_6$. **II,b)** Time-dependent changes in protein concentrations of nodes $N_2$ (red curve) and $N_3$ (blue curve). **II,c)** Protein concentration dynamics of the output node $N_6$. **II,d)** Fourier transform of the output protein concentration (blue curve), with the calculated frequency of the dual-feedback oscillator (red dashed line) and the expected frequency peaks corresponding to the modulation with the CRISPRlator's frequency (black dashed lines). The simulations for the AM lasted for $10^5$ minutes, with the first $10^3$ minutes treated as transient time. In this case an absolute tolerance of $10^{-12}$ and a relative tolerance of $10^{-10}$ was used. We described the dual-feedback oscillator using a slightly modified Elowitz model, where the protein degradation rate was set to $d_P = 0.2$ molecules. To induce oscillations, we introduced higher nonlinearity by using a Hill exponent of 6 for the activation and repression interactions between nodes $N_1$ and $N_6$. The detailed descriptions of the models are available in the SI as "beat.html" and "AM_independent_Stricker.html", respectively.

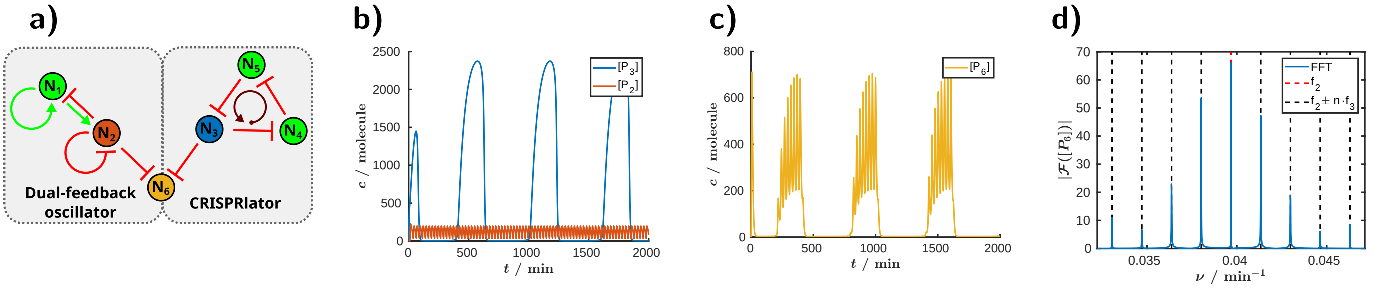

**Figure S5: Effect of complete repression on amplitude modulation.** This figure follows the same panel layout as Figure 3II, but with a complete repression strength from $N_3$ to $N_6$. Under this setting, when $N_2$ and $N_3$ are out of phase, the interaction results in near-complete repression, as shown in panel (c). Detailed model descriptions are provided in the SI file "AM_independent_Stricker.html".

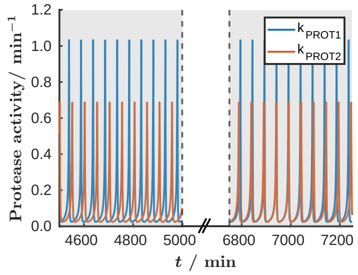

**Figure S6: Temporal changes in protease activity during oscillations.** This figure shows the variation in protease activity corresponding to the simulation in Figure 3b. Although the protease concentration was held constant, its activity changes dynamically according to the Michaelis–Menten equation, due to time-dependent fluctuations in the concentration of its protein substrates. The protease activity was calculated for the first protease ($\mathrm{PROT}_1$) using the following expression: $\mathrm{k}_{\mathrm{prot,max}}\,[\mathrm{PROT}_1]/(\mathrm{K}_{\mathrm{prot}} + [\mathrm{uP}_{\mathrm{N1}}] + [\mathrm{P}_{\mathrm{N1}}] + [\mathrm{uP}_{\mathrm{N2}}] + [\mathrm{P}_{\mathrm{N2}}] + [\mathrm{uP}_{\mathrm{N3}}] + [\mathrm{P}_{\mathrm{N3}}])$, where $\mathrm{k}_{\mathrm{prot,max}}$ is the maximum catalytic rate, $\mathrm{K}_{\mathrm{prot}}$ is the Michaelis constant, and [PROT], [uP], and [P] denote the concentrations of the protease, unfolded protein, and folded protein, respectively, with indices referring to the corresponding node numbers. The detailed descriptions of the models are available in the SI as "beat.html".

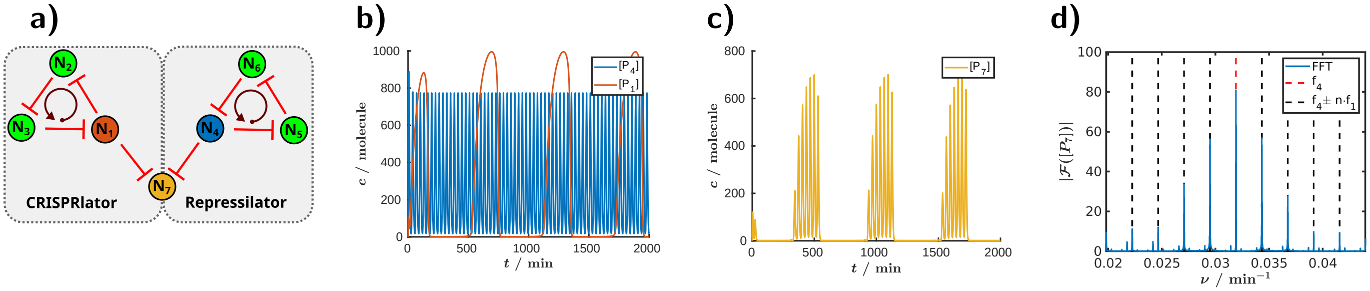

**Figure S7: Amplitude Modulation:** The same configuration as in Figure 3, but with an "accelerated" repressilator coupled to the CRISPRlator. **a)** The topology of the circuits, with a CRISPRlator and a repressilator connected via a NOT gate at node $\mathrm{N}_7$. **b)** Time-dependent changes in protein concentrations of nodes $\mathrm{N}_1$ (red curve) and $\mathrm{N}_4$ (blue curve). **c)** Protein concentration dynamics of the output node $\mathrm{N}_7$. **d)** Fourier transform of the output protein concentration (blue curve), with the calculated frequency of the repressilator (red dashed line) and the expected frequency peaks corresponding to the modulation with the CRISPRlator's frequency (black dashed lines). The simulations for the AM lasted for $10^5$ minutes, with the first $10^3$ minutes treated as transient time. In this case an absolute tolerance of $10^{-14}$ and a relative tolerance of $10^{-12}$ was used and the repressilator was speed up by using a larger degradation rate for the proteins: $d_P = 0.5$ molecules. The time period of the CRISPRlator is 597.2 minutes, while the time period of the repressilator with the increased protein degradation rate is 31.3 minutes. The detailed descriptions of the models are available in the SI as "AM_independent.html".

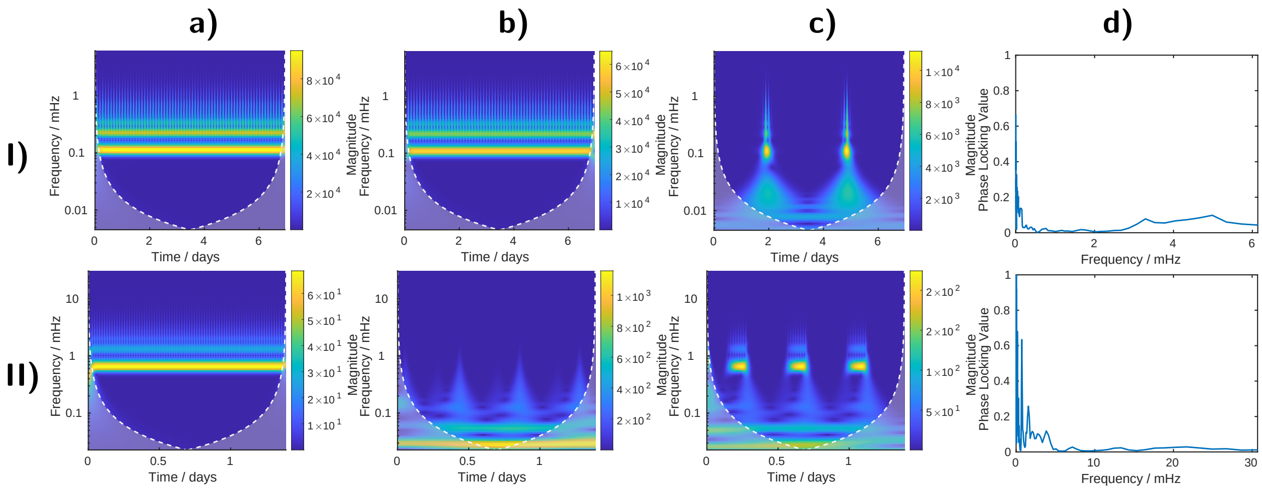

**Figure S8: Wavelet analysis of independently coupled oscillators.** Rows (I) and (II) correspond to simulations of two distinct circuits, as shown in Figure 3. **a–c)** Continuous wavelet transforms of the signals from the two circuits: I,a) $P_{N1}$, I,b) $P_{N4}$, I,c) $P_{N7}$ for the first circuit; II,a) $P_{N2}$, II,b) $P_{N3}$, II,c) $P_{N6}$ for the second. **d)** Phase-locking value (PLV) as a function of frequency, calculated from the phase difference between the CWT representations in panels (a) and (b). Panels (a) and (b) demonstrate that the input signals maintain distinct frequencies, indicating the absence of frequency synchronization. Panel (c) shows the time-resolved frequency spectrum of the output signal. Panel (d) further confirms the lack of phase locking between the oscillators.

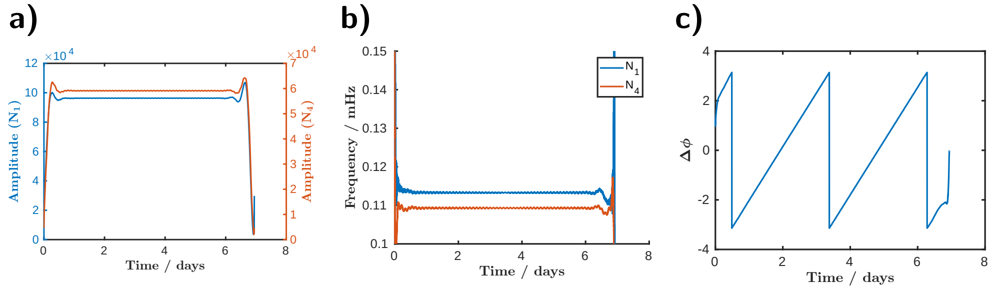

**Figure S9: Analysis of independently coupled oscillators using the Hilbert transform.** Instantaneous amplitude, frequency, and phase difference are shown in panels **a)**, **b)**, and **c)**, respectively, obtained by bandpass filtering and Hilbert transform as described in Section 2.1.4, for the simulation in Figure 3I and Figure 2.1.5I (nodes $N_1$ and $N_4$). In the absence of coupling, both frequency and amplitude remain constant, while the phase difference drifts continuously. For comparison, the corresponding weakly coupled case is shown in Figure 4.

### 3 Deeply coupled oscillators

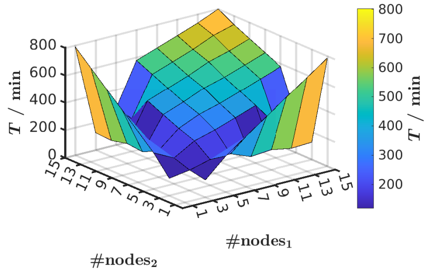

**Figure S10: Three-dimensional version of Figure 4e.** This figure illustrates the linear relationship between the number of nodes and the oscillation period in coupled repressilator systems.

#### 3.1 Alternative strategies for tuning promoter strength in a circuit: adjusting the protease concentration and modifying the repression strength

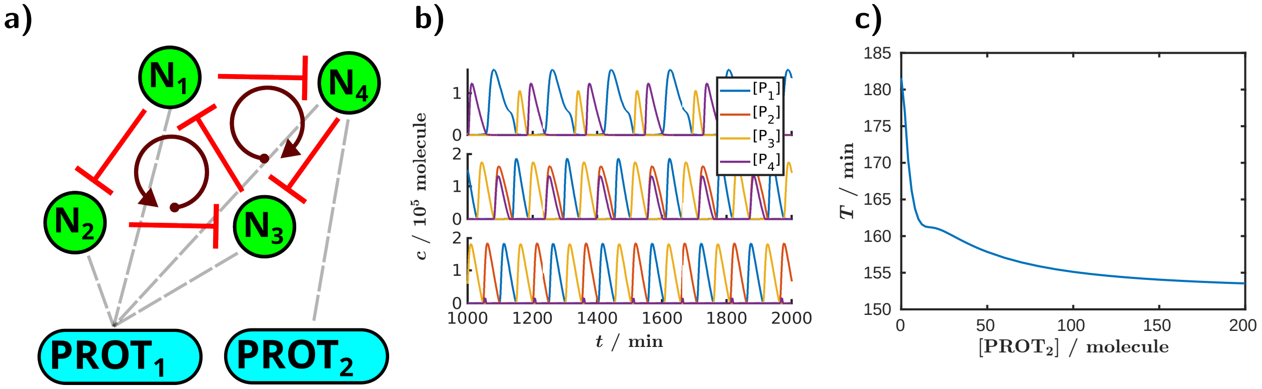

**Figure S11: Deeply coupled oscillators simulated using the Tomazou model [3], analogous to Figure 4.** Figures 4, S11, and S12 demonstrate that the same phenomenon can be observed through multiple approaches: varying the promoter strength (Figure 4), altering the protease concentration (Figure S11), or adjusting the repression strength (Figure S12). $PROT_1$ was held constant at 100 molecules, while the concentration of $PROT_2$ was varied. **a)** Topology of the circuit. **b)** Protein oscillations at different nodes over time with 0, 50, and 500 molecules of $PROT_2$. **c)** Time period changes as a function of $PROT_2$ concentration. The simulations ran for $10^4$ minutes, with the first $10^3$ minutes regarded as transient time. The detailed descriptions of the model for the circuit in a) is available in the SI as "deeply_coupled_Tomazou.html".

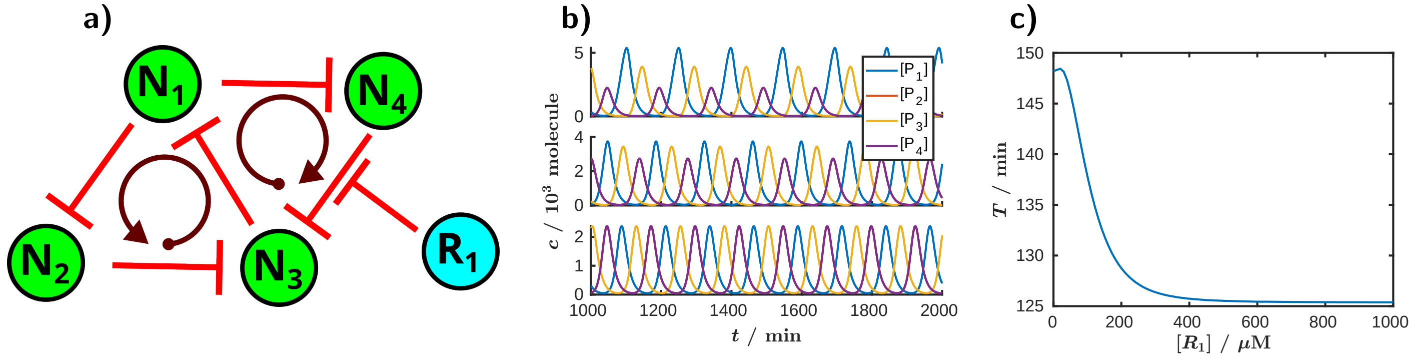

**Figure S12: Deeply coupled oscillators simulated using the Elowitz model [2], analogous to Figure 4.** Figures 4, S11, and S12 demonstrate that the same phenomenon can be observed through multiple approaches: varying the promoter strength (Figure 4), altering the protease concentration (Figure S11), or adjusting the repression strength (Figure S12). In this case, the promoter strength was held constant, while the concentration of the inducer $R_1$ was varied. **a)** Topology of the circuit. **b)** Protein oscillations at different nodes over time with 0, 100, and 1000 $\mu$M of $R_1$. In this case, nodes $N_2$ and $N_4$ are equivalent because they are both repressed solely by node $N_1$, leading to identical protein oscillations. As a result, the curve for N2 is not visible in the graph, as it overlaps with the curve for $N_4$. **c)** Time period changes as a function of $R_1$ concentration. The simulations ran for $10^4$ minutes, with the first $10^3$ minutes regarded as transient time. The detailed descriptions of the model for the circuit in a) is available in the SI as "deeply_coupled_Elowitz_inducer.html".

#### 3.2 The deeply coupled Goodwin oscillator

In this section, we first examine the properties of the Goodwin oscillator and then demonstrate that its characteristic oscillatory behavior is not limited to a single self-repressing node, it can also emerge in analogous multi-node systems. Notably, we show that even the repressilator, despite being known for its sequential oscillations of the three nodes, can exhibit Goodwin-type behavior under specific conditions. The Goodwin oscillator consists of a single node that represses itself [5–8]. Early models often implemented high non-linearity and delayed negative feedback to induce oscillations [9, 10]. Although these models successfully captured basic oscillatory behavior, the nonlinearity in the applied Hill function was unrealistically high, indicating the presence of additional hidden reaction steps. Later, Bharath *et al.* demonstrated that this high Hill coefficient could be avoided by incorporating self-activating positive feedback or using Michaelis-Menten degradation kinetics [11, 12].

Oscillators in chemistry and biology generally rely on two essential components: auto-activation and delayed negative feedback [13]. The Goodwin oscillator contains these essential components; indeed, in Figure S13a, we illustrate how the Goodwin oscillator is qualitatively analogous to such a system when applied to a protease-based model like the one described by Tomazou [3], which we use in our simulations. The middle topology in Figure S13a displays that the amount of folded protein (P) increases with the amount of unfolded protein (uP). Due to the negative feedback in the self-repressing node, the folded protein represses the unfolded protein (by binding to the promoter and reducing mRNA synthesis), which subsequently lowers uP levels with a delay. The system also includes an implicit auto-activation step (through the protease system), which is beneficial for creating robust oscillations. The protease degrades protein-based transcription factors (represented by a repression symbol in the figure), but when these transcription factor levels are sufficiently high, they saturate the proteases, reducing their relative activity. This is symbolized in the figure as repression of the protease's activity. By slowing their own degradation, transcription factors can indirectly increase their own concentration, represented as auto-activation in the diagram. As a result, the system features both auto-activation and delayed negative feedback, often leading to stable oscillations. The bifurcation diagram shows that the oscillations are generated by a supercritical Hopf bifurcation (Figure S13b), while Figure S13c illustrates a wide oscillatory region as a function of protein maturation time (delay) and protease concentration. A wide oscillation region is crucial to increase the chances of observing oscillations in an experimental implementation.

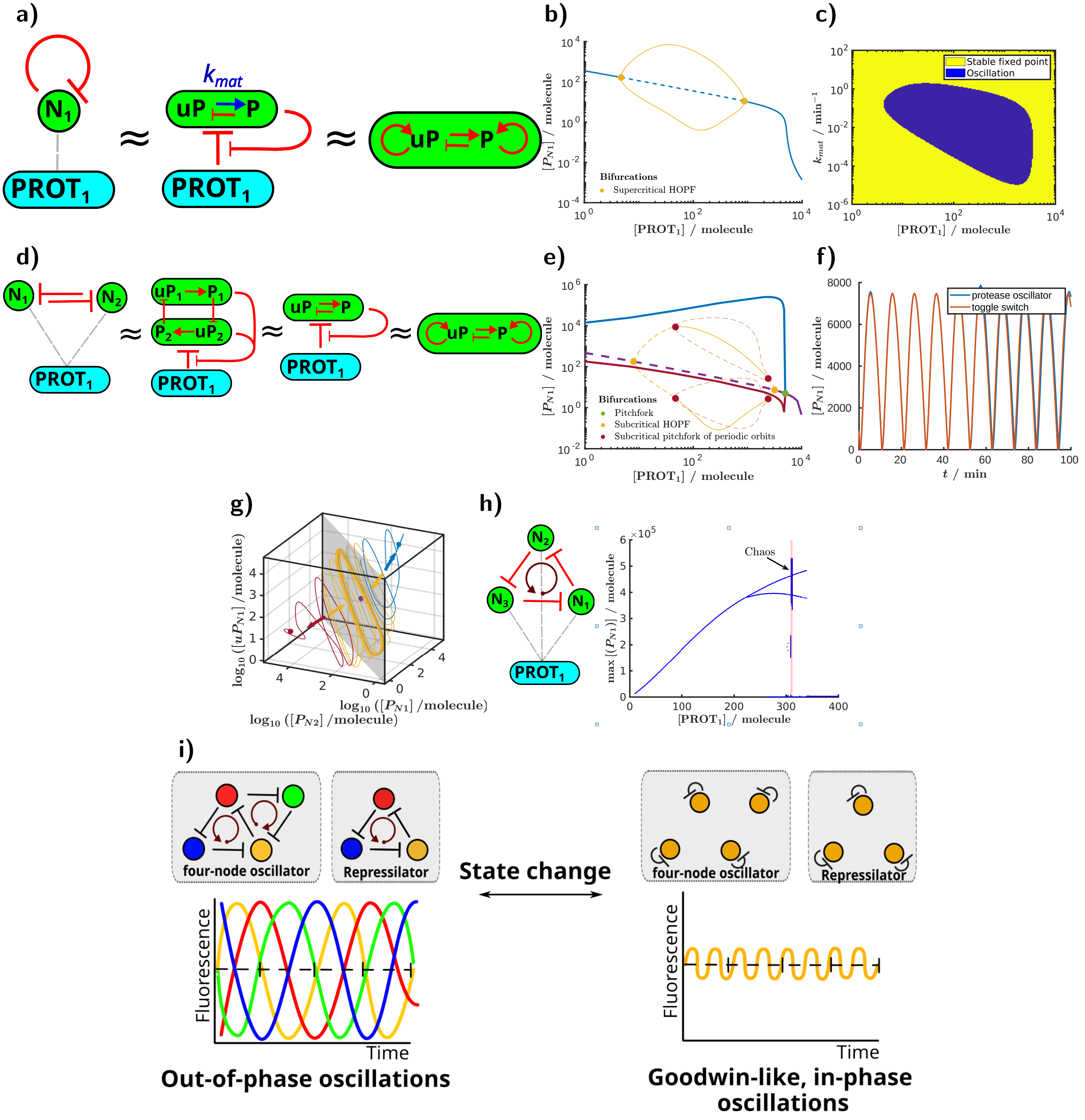
a)
$k_{mat}$
uP
P
PROT₁
b)
Bifurcations
Supercritical HOPF
[PROT₁] / molecule
$[P_{N1}]$ / molecule
c)
Stable fixed point
Oscillation
$k_{mat}$ / min⁻¹
[PROT₁] / molecule
d)
e)
Bifurcations
Pitchfork
Subcritical HOPF
Subcritical pitchfork of periodic orbits
f)
protease oscillator
toggle switch
t / min
g)
h)
Chaos
[PROT₁] / molecule
i)
four-node oscillator
Repressilator
State change
Fluorescence
Time
Out-of-phase oscillations
Goodwin-like, in-phase oscillations

**Figure S13: The Goodwin oscillator and analogous oscillations in other circuits.**
**a)** Topology of the Goodwin oscillator in different representations. $PROT_1$ represents the protease, while $N_1$ denotes the node where the unfolded protein-based transcription factor (uP) matures into a folded protein (P) at a maturation rate of $k_{mat}$. **b)** Bifurcation diagram of the Goodwin oscillator showing protein concentration N1 as a function of protease concentration $PROT_1$. Blue line: stable fixed points; dashed blue line: unstable fixed points; continuous yellow line: stable periodic orbits (upper and lower bound of the limit cycle). The yellow dots sign the supercritical Hopf bifurcations. **c)** The oscillatory region (blue) of the Goodwin oscillator as a function of protease concentration and maturation rate, $k_{mat}$. The $PROT_1$ protease concentration was set to 50 molecules. **d)** Topology of the toggle switch in different representations. **e)** Bifurcation diagram of the toggle switch showing the protein concentration of the first node ($N_1$) as a function of protease concentration ($PROT_1$). Blue, red, and purple lines: stable fixed points; dashed yellow and orange lines: unstable periodic orbits; continuous yellow lines: stable periodic orbits (upper and lower bound of the limit cycle). The green dot indicates the pitchfork bifurcation, while the yellow dots mark the subcritical Hopf bifurcation of fixed points. The red dots represent the subcritical pitchfork bifurcations of periodic orbits. **f)** Trajectory of protein concentration over time for the Goodwin oscillator (blue) and the toggle switch in its oscillatory state (red). For the Goodwin oscillator, we used $K_{protease} = 15$ molecules and 50 molecules of protease, which is half the value applied in the toggle switch. **g)** Phase space representation of the toggle switch. The red and blue dots represent stable fixed points, while the purple dot indicates an unstable fixed point. The stable limit cycle (orange) is shown on the gray plane, where the two nodes are synchronized (the concentrations of corresponding species in the nodes are equal). Four trajectories are depicted in the phase space, color-coded according to the attractor they approach: red and blue for the stable fixed points, and orange for the limit cycle. The protease concentration was set to 100 molecules. **h)** Topology and bifurcation diagram for the repressilator. The blue curve represents the maximum protein concentration for node $N_1$, while the red region highlights where the largest Lyapunov exponent is positive. Chaotic oscillators are characterized by positive Lyapunov exponents. **i)** Comparison of out-of-phase oscillations and Goodwin-type in-phase oscillations in multi-node circuits. Further simulations for specific protease concentrations can be found in Figure S16. Additional simulation details can be found in the Methods section and the SI, specifically in the files "goodwin.html" and in "repressilator_Tomazou.html".

Intriguingly, we predict that the toggle switch [14–16], a well-studied network of two mutually repressive nodes, can exhibit a similar Goodwin-type oscillatory state (Figure S13d). However, in this design, a protease and intermediate unfolded proteins are essential components of the circuit. To emphasize their central role, we therefore refer to this circuit as the oscillatory toggle switch. The transcription factors $P_1$ and $P_2$ in the oscillatory toggle switch repress the production of the unfolded protein in the opposite node ($uP_2$ and $uP_1$, respectively). The subsequent steps, involving the proteases and the production of folded proteins, are analogous to those in the Goodwin oscillator. For the oscillatory toggle switch to exhibit the same self-accelerating behavior as the Goodwin oscillator, the proteins must synchronously increase and decrease their concentrations. In other words, for the system to undergo Goodwin-type oscillations, both nodes must oscillate in synchrony. To symbolize this synchronization, we replace $P_1$ and $P_2$ with a single notation, P, and similarly, we use uP instead of $uP_1$ and $uP_2$. As demonstrated in the final step of Figure S13d, once the nodes are synchronized, the oscillatory toggle switch qualitatively exhibits a behavior analogous to the oscillator observed in the last step of Figure S13a with the Goodwin oscillator. However, Figure S13d presents only a logical simplification of the underlying mechanism to help illustrate how this circuit functionally corresponds to a system with self-activation and delayed negative feedback, a well-known motif capable of generating oscillations.

These qualitative observations raise an important question: is this oscillatory state realistic, and how could it be achieved experimentally? To explore this, we analyzed the system's bifurcation diagram (Figure S13e). The diagram reveals three distinct regimes depending on the protease concentration. At high protease concentrations, the system exhibits a single stable fixed point characterized by low protein levels (solid purple line). At low and high protease levels (excluding the intermediate range) the system is bistable and behaves like a classic toggle switch: one node is activated (solid blue), the other deactivated (solid red), with an unstable intermediate state (dashed purple). However, within an intermediate protease concentration range, two stable periodic orbits emerge (solid orange lines), corresponding to fast, low-amplitude in-phase oscillations similar to those seen in the Goodwin oscillator. In this regime, the system becomes multistable. Depending on the initial conditions, it can exhibit either the traditional toggle switch behavior or settle into the oscillatory state. More specifically, when the system is initialized asymmetrically, with one node significantly more active (higher protein and mRNA levels), it behaves like a toggle switch. In contrast, when both nodes start with similar activation levels, the system tends to evolve toward the Goodwin-type oscillatory behavior. In other words, the toggle switch and oscillatory states can coexist, and the observed dynamics are determined by the initial symmetry of the system. A detailed analysis of the bifurcation structure is provided in the SI, Section 3.2.2.

We further asked ourselves whether this resemblance to the Goodwin oscillator is merely qualitative, or is there a quantitative agreement? In Figure S13f, we present a simulation comparing the Goodwin oscillator and the oscillatory toggle switch. The results show that when the nodes of the toggle switch are synchronized (in-phase), the oscillations of the Goodwin oscillator and the oscillatory toggle switch are in complete agreement and the system quantitatively exhibits Goodwin-type oscillations. In other words, when the nodes are synchronized – meaning the concentrations of the appropriate species (produced by the nodes) are equal for both nodes – the system resides in a subspace of the phase space of the oscillatory toggle switch that corresponds directly to the Goodwin oscillator. Within this subspace, the fixed point and periodic orbit are the same as those found in the Goodwin oscillator (Figure S13g, gray region). However, the trajectories can leave this subspace and move toward the stable fixed points of the oscillatory toggle switch (red and blue arrows), making the fixed points and periodic orbits unstable within the subspace. Interestingly, we observe a region where the periodic orbit remains stable (orange arrow), and as demonstrated in the simulations, this stable orbit matches the behavior of the Goodwin oscillator. In summary, our simulations indicate that when the two nodes of the oscillatory toggle switch begin from similar initial states, the protease-mediated coupling can synchronize them. As a result, the circuit behaves like a single self-repressing node, capable of exhibiting oscillatory behavior (Figure S13d–g). To the best of our knowledge, toggle switch oscillations have not yet been demonstrated experimentally, making these findings novel theoretical predictions.

Similarly, we theoretically predict that the repressilator family (Figure S13h) could also establish synchronized, in-phase, Goodwin-type oscillations. In this case, we observe deeply coupled oscillators, where the coexistence of two oscillatory states – asynchronous (well-known, the three nodes oscillate out-of-phase) and synchronous (Goodwin-type, the three nodes oscillate in-phase) – leads to the emergence of new and interesting dynamical phenomena (Figure S13i). We initialized the repressilator in an asymmetric state, where initial protein concentrations differed between nodes, and analyzed its bifurcation diagram (Figure S13h). The multistable system switches from the out-of-phase oscillations to the low-amplitude, high-frequency oscillatory state at higher protease concentrations. The time period of the out-of-phase oscillations is approximately 11 times longer, while their amplitude is roughly 19 times greater compared to the in-phase oscillations. Starting from a symmetric initial condition, it is possible to achieve the Goodwin-type oscillatory state, even at lower protease concentrations (Figure S15). There is also an intermediate region where coupling effects between different oscillatory states lead to period-2 oscillations and even chaotic behavior in a small range. It is fascinating to observe, even theoretically, such exotic dynamical behavior in one of the most well-known oscillators in synthetic biology. Although this chaotic region is too narrow to be experimentally accessible, it suggests that other parameter sets or circuits could be explored to create an autonomous chaotic oscillator. For example, deeply coupling two repressilators ($N_1$-$N_2$-$N_3$ and $N_1$-$N_4$-$N_3$) to each other leads to a significantly broader chaotic region (Figure S14), making the experimental realization of chaotic oscillations more feasible. Beyond this coupling, interactions between in-phase and out-of-phase oscillations contribute to complex dynamical behavior. Although a detailed analysis of this system requires a separate study, it is evident that the interaction between the two repressilator rings further destabilizes the system, thereby expanding the chaotic region.

**Finding unstable periodic orbits** As discussed in the Results section, the Goodwin oscillator corresponds to a subspace of the toggle switch, where the concentrations of the two nodes are equal. Using this insight, we located the periodic orbit in this subspace by applying the Goodwin oscillator, assuming that the amount of protease enzymes and the Michaelis constant ($K_{protease}$) is halved in the Goodwin oscillator compared to the toggle switch. However, trajectories can leave this subspace toward stable fixed points, destabilizing a part of the periodic orbit. We observed that this unstable periodic orbit (UPO) stabilizes through a pitchfork bifurcation of periodic orbits, which simultaneously creates two new UPOs. We identified these UPOs as follows: first, we perturbed the system around the bifurcation point where the UPOs are close to the stable periodic orbit (SPO). We calculated the distance in phase space between the initial and final states after an estimated time period. Then, we minimized this distance by adjusting the initial concentrations and time period using MATLAB's `fminsearch` and `lsqnonlin` functions, setting both `FunctionTolerance` and `StepTolerance` to $10^{-12}$ for high accuracy. Once we identified the starting point of the UPO, we applied a continuation method by gradually altering the protease concentration in small steps (1 molecule per step), repeating the minimization process to track the UPO. If the minimum was not found, we reduced the step size in the protease concentration by half. After determining the initial concentrations and time period for the UPO, we calculated the maximum and minimum protein values during the first cycle to generate the bifurcation diagram. Since the system is symmetric, finding one UPO suffices, as the second UPO corresponds to the situation where the two nodes are swapped.

###### 3.2.1 Fixed Point and Linear Stability Analysis

The dynamical system under study is represented by a set of $n$ coupled nonlinear ordinary differential equations (ODEs), derived from the underlying biochemical reaction network:

$$\frac{\mathrm{d}\mathbf{x}}{\mathrm{d}t} = \mathbf{f}(\mathbf{x}, \mu), \tag{S26}$$

where $\mathbf{x} \in \mathbb{R}^n$ denotes the vector of biochemical species concentrations, and $\mu \in \mathbb{R}$ is a key control parameter (e.g., protease concentration) acting as the bifurcation parameter.

**Fixed Points.** The steady states (fixed points) of the system, $\mathbf{x}^*$, satisfy

$$\mathbf{f}(\mathbf{x}^*, \mu) = \mathbf{0}. \tag{S27}$$

For each value of the bifurcation parameter $\mu$, Equation (S27) was solved numerically to identify all admissible steady states.

**Linear Stability Analysis.** To determine the stability of each fixed point $\mathbf{x}^*$, we linearized the dynamics in its vicinity. Small perturbations $\delta\mathbf{x} = \mathbf{x} - \mathbf{x}^*$ evolve according to the Jacobian matrix $\mathbf{J}$:

$$\frac{\mathrm{d}}{\mathrm{d}t}(\delta\mathbf{x}) \approx \mathbf{J}(\mathbf{x}^*)\,\delta\mathbf{x}, \qquad J_{ij} = \left.\frac{\partial f_i}{\partial x_j}\right|_{\mathbf{x}^*}. \tag{S28}$$

The eigenvalues $\lambda$ of $\mathbf{J}(\mathbf{x}^*)$ determine local stability. The classification of fixed points, together with the identification of stable and unstable periodic orbits, is described in detail in the Methods section of the main text.

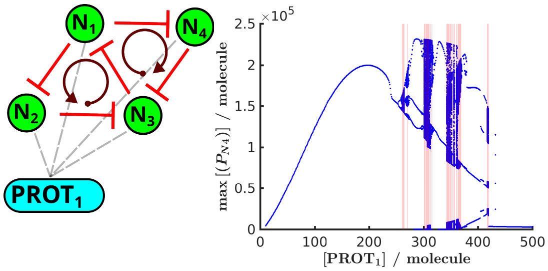

**Figure S14: Topology and bifurcation diagram for the deeply coupled repressilator.** The blue curve represents the maximum protein concentration for node $N_1$, while the red region highlights where the largest Lyapunov exponent is positive. Chaotic oscillators are characterized by positive Lyapunov exponents. Further simulations for specific protease concentrations can be found in Figure S16. Additional simulation details can be found in the Methods section and the SI, specifically in the file "deeply_coupled_Tomazou.html".

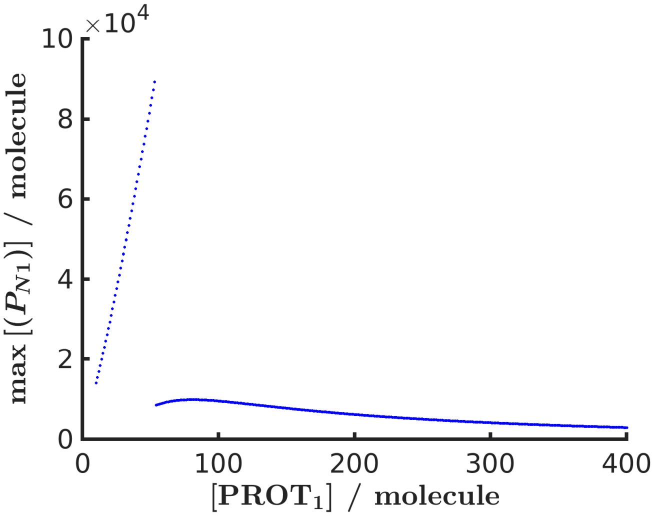

**Figure S15: Bifurcation diagram for the repressilator, showing the maximum protein oscillation for the first node as a function of protease concentration.** The simulation started from a symmetric initial condition, resulting in a Goodwin-type oscillatory state characterized by low-amplitude, fast oscillations. For comparison, Figure S13h shows the bifurcation diagram obtained with asymmetric initial conditions.

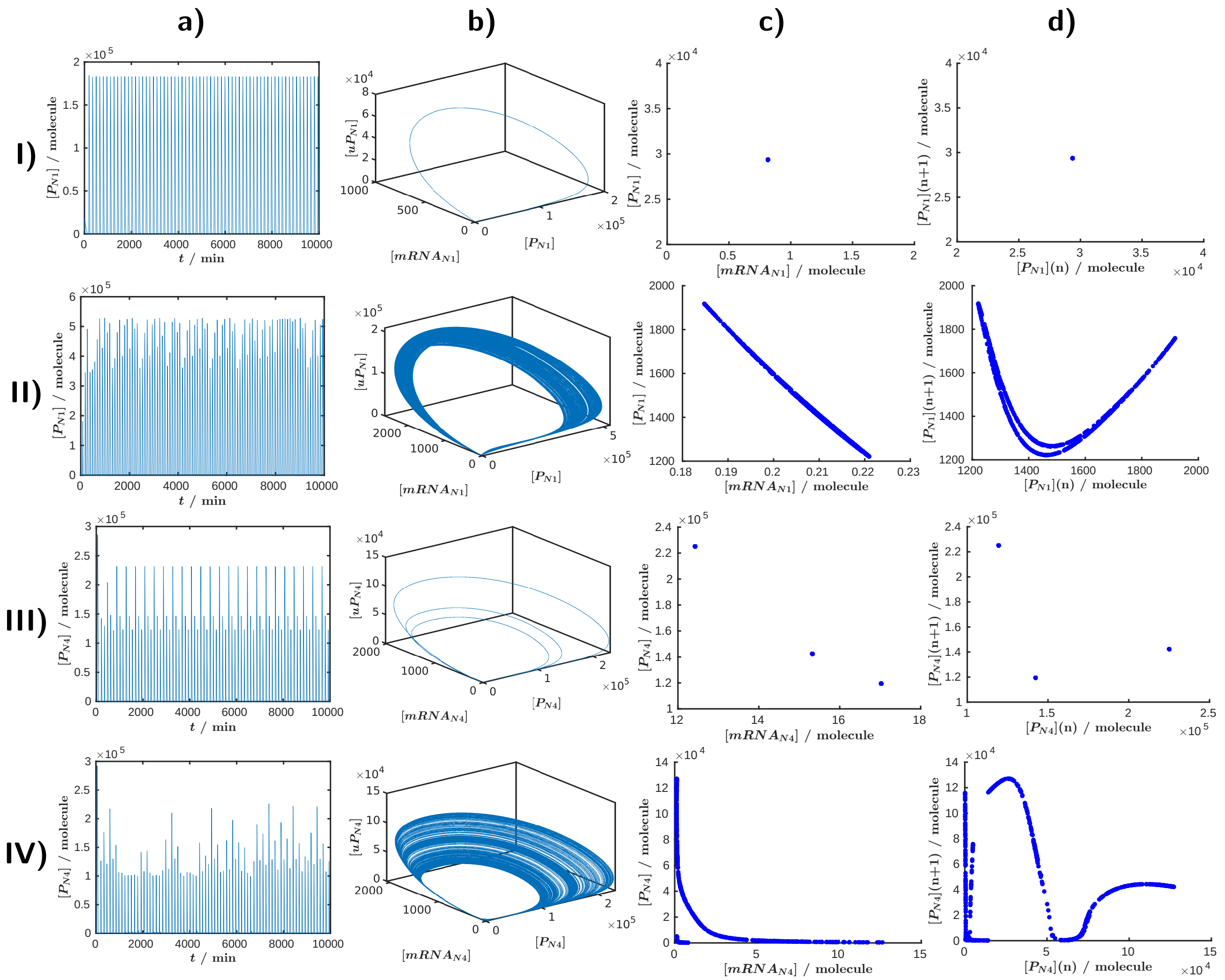

**Figure S16: Additional simulations for deeply coupled oscillators, as shown in Figure S13h and in Figure S14.**
**Rows: I)-II)** Repressilator with protease concentrations of $\mathrm{PROT}_1 = 100$ molecules (I) and $\mathrm{PROT}_1 = 310$ molecules (II). **III)-IV)** deeply coupled repressilator with protease concentrations in the second circuit of $\mathrm{PROT}_2 = 290$ molecules (III) and $\mathrm{PROT}_2 = 310$ molecules (IV).
**Columns: a)** Protein concentration in the first node over time. **b)** Phase space diagram showing the unfolded protein-based transcription factor (uP), mRNA, and folded protein (P) concentrations for the first node in the repressilator and the fourth node in the deeply coupled repressilators. **c)** Poincaré section at $10^2$ molecules of uP concentration (crossing from the positive direction). **d)** Poincaré map derived from the Poincaré section in (c).

##### 3.2.2 Comprehensive description of the bifurcation behavior in the toggle switch

Figure S13e shows the bifurcation diagram as a function of protease concentration, providing insight into the system's behavior relative to protease levels. This allows us to determine whether the oscillatory state indeed arises in this system. At high protease levels, there is only one stable (i.e. non-oscillating) state with low transcription factor concentration (purple curve). As the protease concentration decreases, a pitchfork bifurcation occurs, leading to two stable states (red and blue) and one unstable state (dashed purple). This is the typical behavior of the toggle switch, where one node exhibits high protein concentration and the other exhibits low concentration, while the intermediate state is unstable. If we further decrease the protease concentration, first we can observe a subcritical Hopf bifurcation, where an unstable periodic orbit (UPO) emerges (dashed orange) and this orbit becomes stable in a subcritical pitchfork bifurcation of periodic orbits [17] (continuous orange), and two UPOs are created (dashed red). The pitchfork bifurcation is common in symmetrical systems, hence its multiple appearances in our symmetric toggle-switch model. Interestingly, across a broad range of protease concentrations, we can observe a stable periodic orbit (SPO, continuous orange), where the toggle switch enters a Goodwin-type oscillatory state with synchronized nodes. In this region the system shows multistability, thus we can observe both a SPO and two stable fixed points.

### 4 Weakly and strongly coupled oscillators

**Extended caption for Figure 5**: **Synchronization, frequency and amplitude modulation, chaos and resonance with coupled oscillators.**
**a)-d) Synchronization between coupled repressilators.** **a)** Topology, where two oscillators are coupled through a shared protease, $\mathrm{PROT}_3$. **b)** Phase difference over time ($\Delta t$) for various $\mathrm{PROT}_3$ protease concentrations (1, 2, 4, 8, and 16 molecules) (top) and protein oscillations in the $\mathrm{N}_1$ node with slightly different initial conditions for the two oscillators: $\mathrm{P}_{\mathrm{N1}}]_0 = 1000$ molecules, while $\mathrm{P}_{\mathrm{N4}}]_0 = 600$ molecules (bottom). **c)** Protein trajectories in the $\mathrm{N}_1$ node for 0, 40, and 100 molecules of $\mathrm{PROT}_3$, respectively. **d)** Variation in the average time period (T) of the first (blue) and fourth (red) nodes as a function of the common protease ($\mathrm{PROT}_3$)concentration. In c) and d) the two repressilators have a different frequencies due to the different amount of proteases: $\mathrm{PROT}_1 = 50$ molecules, $\mathrm{PROT}_2 = 200$ molecules.
**e)-h) Higer period oscillations and chaos.** **e)** Topology of two unidirectionally coupled repressilators. **f)** Bifurcation diagram for the circuits in e) Additional simulations for this circuit are presented in Figure S20I,II. **g)** Topology of a Goodwin oscillator unidirectionally coupled to a repressilator. **h)** Bifurcation diagram of the circuits in g). In the bifurcation diagrams (f and h), the maximum of the protein oscillations in the first node is plotted as a function of the protease concentration in the driving system, $\mathrm{PROT}_2$. The red colour indicates regions of chaos.
**i)-k) Resonance.** Additional simulations for this circuit are presented in Figure S20C,D. **i)** Topology of two unidirectionally coupled repressilators. **j)** Changes in the time period of the second oscillator as a function of the promoter strength of the sixth node. **k)** Amplitude of the oscillations at the $\mathrm{N}_1$ node as a function of the promoter strength of the $\mathrm{N}_6$ node, $k_{1,N6}$. The simulations were $2 \cdot 10^4$ minutes long, with the half part considered as transient. For the resonance-based simulation (i)-k)), we employed the Elowitz model [2, 18], while for the other cases (a)-h)), we used the Tomazou model [3]. The details of the simulations can be found in the SI, specifically in "coupled_repressilators_Tomazou.html", "repressilator_repressilator.html", "repressilator_goodwin.html", "resonance.html", respectively.

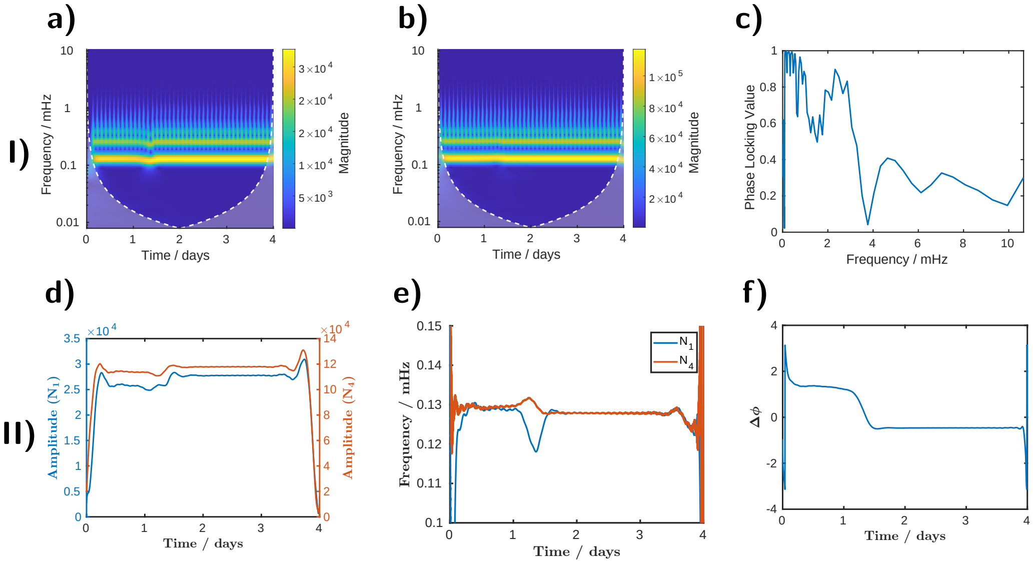

**Figure S17: I): Wavelet analysis of weakly coupled oscillators.** The analysis was performed on the simulation shown in Figure 5d, using 60 molecules of $PROT_3$. **a,b)** Continuous wavelet transforms of the protein concentrations of $P_{N1}$ and $P_{N4}$, respectively. **c)** Phase-locking value calculated from the phase difference between the CWT representations in panels (a) and (b), evaluated after 3 days of simulation. The elevated PLV at low frequencies indicates sustained phase-locking, while differences at higher frequencies arise from shape differences between the signals. The perturbation observed in the main frequency of panel (a) around 1.5 days reflects synchronization, in contrast to the independent oscillator case, where frequencies remain unchanged. **II) Analysis of weakly coupled oscillators using the Hilbert transform.** Instantaneous amplitude, frequency, and phase difference are shown in panels **d)**, **e)**, and **f)**, respectively, obtained by bandpass filtering and Hilbert transform as described in Section 2.1.4. In panel **f**, the phase difference abruptly converges to zero, marking the onset of complete synchronization. Following this transition, the amplitude exhibits a slight increase (panel **d**), while the oscillator frequencies become fully matched (panel **e**). A calculated phase-locking value of 0.95 indicates that the oscillators exhibit strong, essentially complete synchronization.

#### 4.1 Unidirectional coupling

##### 4.1.1 Frequency and amplitude modulation

With the independent coupled oscillators, we demonstrated how to create a pure AM. When the frequency of the oscillation is constant, just the amplitude is changing (Figure 3II). By unidirectionally coupling two oscillators, we can observe both frequency modulation (FM) and amplitude modulation (AM). In the general scenario illustrated in Figure S18, a slower oscillator (the repressilator) modulates a faster oscillator (the Goodwin oscillator). This approach allows us to connect any faster and slower oscillators (e.g. also the repressilator to the CRISPRlator). However, modifying the promoter strength of a node impacts both the amplitude and frequency of the oscillator, causing changes in amplitude to coincide with frequency shifts. This coupling results in imperfect FM, making it impossible to observe pure AM or FM signals in this system.

To make AM and FM more independent, we use an output node, G, which maintains a consistent amplitude while allowing frequency changes in the oscillator (achieved by modifying the promoter strength of one of the nodes). In this setup, we modulated the frequency of the repressilator using a slower oscillator. This slower oscillator was generated by independently coupling two Goodwin oscillators and leveraging the beat phenomenon to create a slower frequency, resulting in the FM signal. Similarly, we achieved AM with coupled oscillators by varying the strength of the output node. As shown in Figure S19c, this approach is still not perfect; although the amplitude does exhibit slight fluctuations during FM, the changes in frequency are significantly more pronounced.

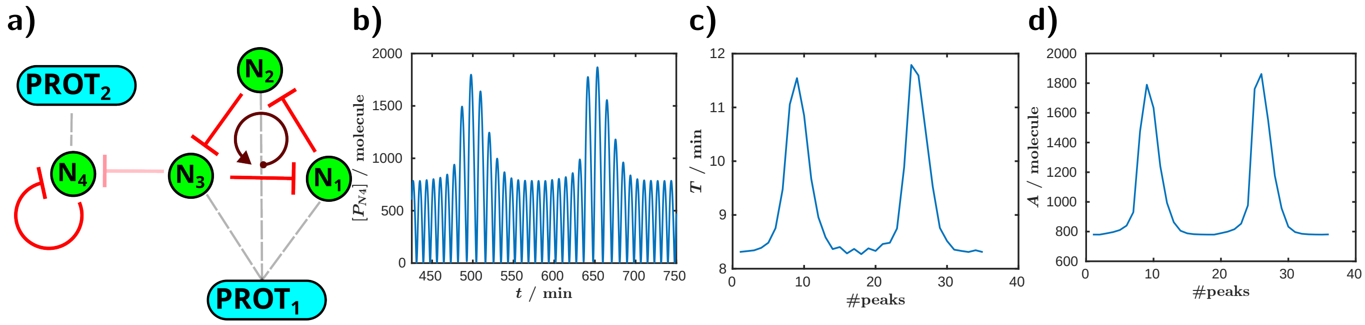

**Figure S18: Not independent frequency and amplitude modulation.** **a)** Topology of the circuit. The slower oscillator (repressilator) drives the faster oscillator (Goodwin oscillator). **b)** Protein oscillations at the fourth node as a function of time. **c) and d)** Variations in the oscillation period and amplitude as a function of the number of peaks. In the frequency modulation we used 50 molecules for half saturation constants in the Hill-function and a weaker repression between the oscillators: $\text{Hill}' = 0.9\text{Hill} + 0.1$. The details of the simulation can be found in the SI as "FM_AM.html".

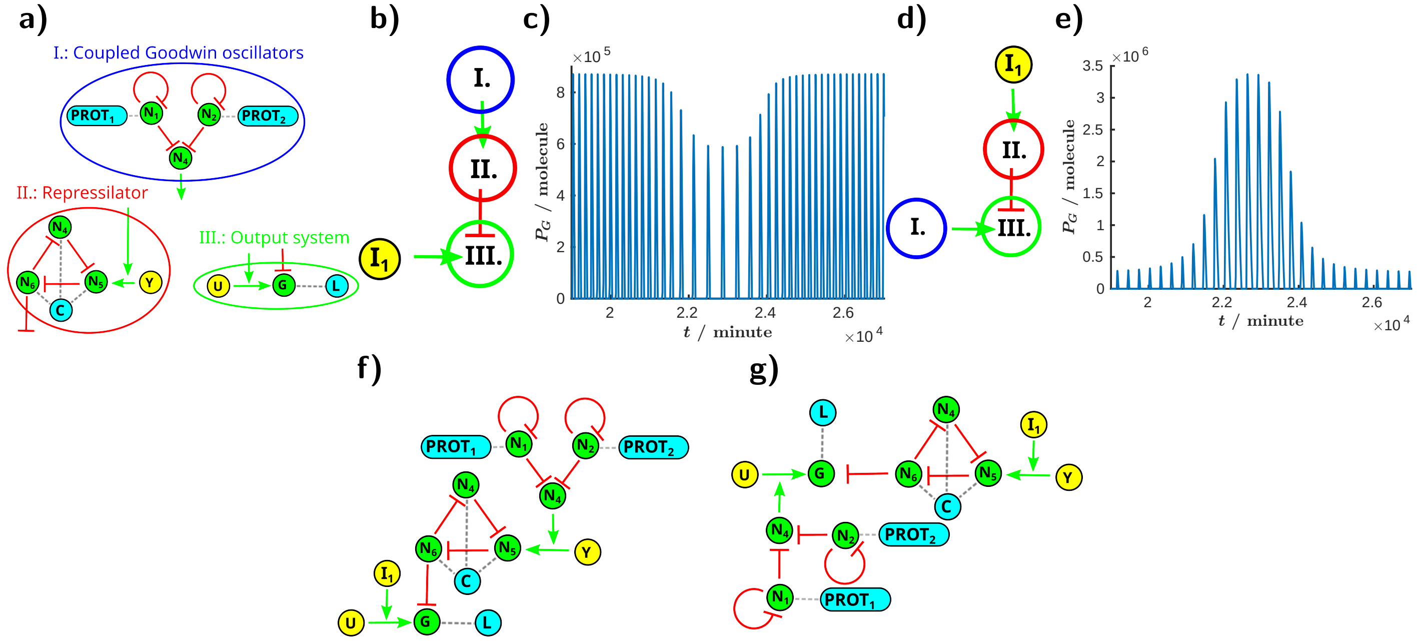

**Figure S19: Frequency and amplitude modulation with intracellularly coupled oscillators. a)** Building blocks for AM and FM in (f) and (h). $PROT_1$, $PROT_2$, C and L represent proteases. U and Y are transcription factors activating G and $N_5$, with $I_1$ functioning as an external inducer. **b)** Simplified representation of the circuit for FM. The coupled Goodwin oscillators (I) modulate the frequency of the repressilator (II), with the output observed in (III). The detailed topology can be found in (j). **c)** Frequency modulation for oscillations at the output node (G) in the topology of e) as a function of time. The concentrations of $PROT_1$ and $PROT_2$ are set to 10 molecules, while the values of $a_1$ in $N_1$ and $N_2$ are 100 and 95 1/minute, respectively. The concentration of $[I_1]$ is 30 $\mu$M, and the half-saturation constant in the Hill function for activating the output of the coupled Goodwin oscillator system ($K$ for $N_3 \rightarrow Y \rightarrow N_5$) is set to 1000 molecules. All other parameters were taken from [3]. **d)** Simplified representation of the circuit for AM. The coupled Goodwin oscillators (I) modulate the amplitude of the output observed in (III), while the repressilator (II) oscillates at a constant frequency. The detailed topology can be found in (k). **e)** Amplitude modulation for oscillations in the topology of g) at the output node (G) as a function of time. All parameters are identical to those in (f), except for the half-saturation constant in the Hill function for activating the output of the coupled Goodwin oscillator system ($K$ for $N_3 \rightarrow U \rightarrow G$), which is set to 500 molecules. **f)** Detailed circuit topology for frequency modulation. **g)** Detailed circuit topology for amplitude modulation. The simulation details for the independent frequency and amplitude modulation can be found in the SI as "FM.html" and "AM.html", respectively.

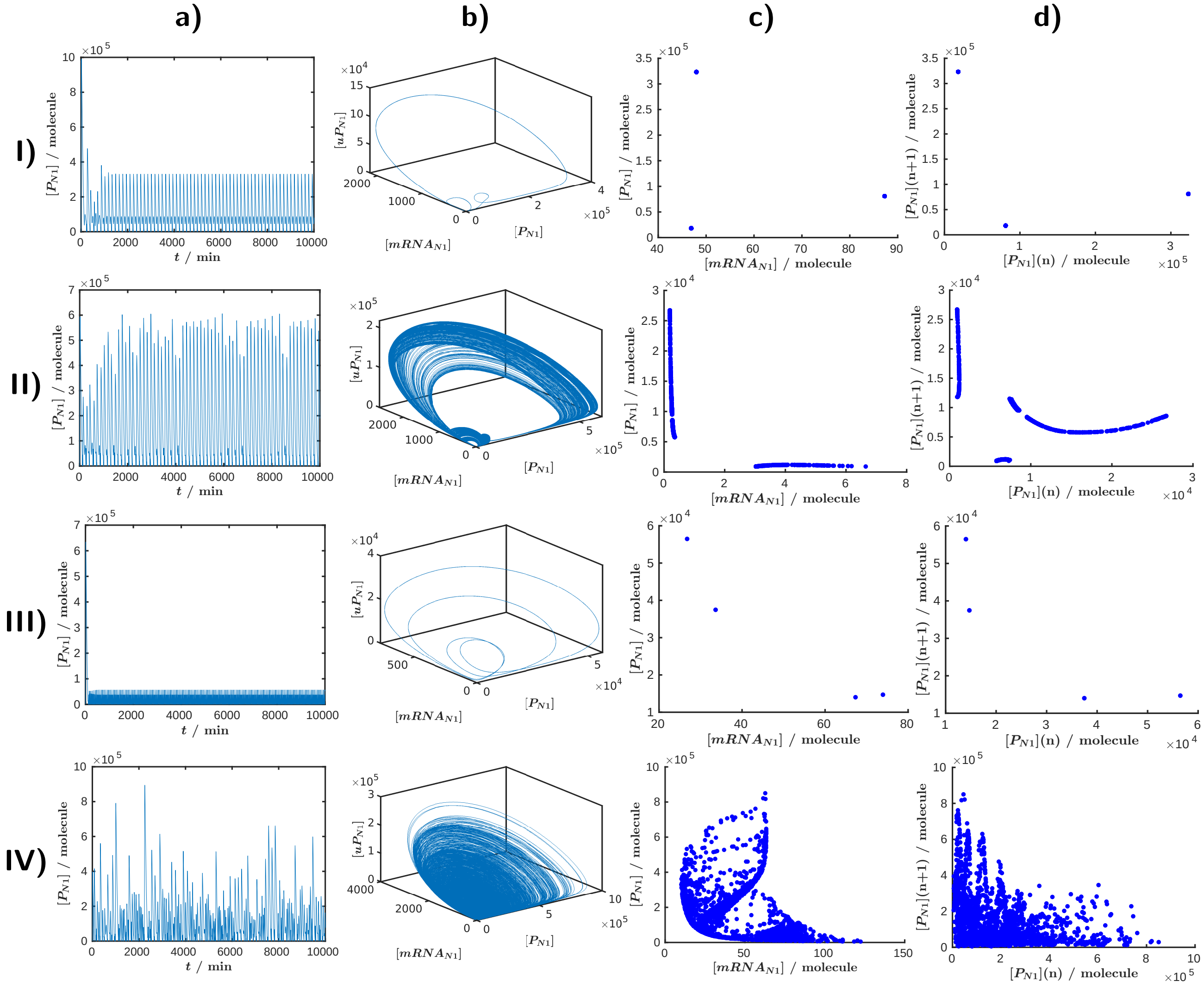

**Figure S20: Additional simulations for unidirectional coupling, as shown in Figure 5.**

**Rows: I)-II)** Unidirectionally coupled repressilators with protease concentrations in the driving system of $PROT_2 = 40$ molecules (A) and $PROT_2 = 150$ molecules (II). **III)-IV)** Unidirectionally coupled Goodwin oscillator and repressilator, with protease concentrations in the Goodwin oscillator of $PROT_2 = 3$ molecules (III) and $PROT_2 = 10$ molecules (IV) (while $PROT_1 = 100$ molecules in the repressilator).

**Columns: a)** Protein concentration in the first node over time. **b)** Phase space diagram showing the unfolded protein ($uP_{N1}$), mRNA ($mRNA_{N1}$), and folded protein ($P_{N1}$) concentrations for the first node. **c)** The Poincaré section is taken at a concentration of $10^2$ molecules in II) and at $10^4$ molecules in all other cases, based on the $uP_{N1}$ concentration (crossing from the positive direction). **d)** Poincaré map derived from the Poincaré section in (c).

### 5 An Application: Oscillator-Based Computers

**Extended caption for Figure 6: Oscillator-Based Computing: Demonstration of Phase Locking with Coupled CRISPRlators.** **a)** Schematic representation of intracellularly coupled oscillator-based computers. We can use small chemical inputs to drive the system and observe a binary output determined by the phase difference between different nodes. **b)** Simplified correlation matrix illustrating in-phase and anti-phase signals, along with an example application of the system operating as an AND gate. **(c, e, g, i) Top row**: Circuit topology, indicating the promoter strength ($a_1$) with values different from unity. The CRISPRlators are coupled via a common dCas system at a low concentration (200 molecules) to enhance coupling strength between the oscillators. **Bottom row**: Protein oscillations for nodes $N_1$, $N_4$, $N_5$, and $N_6$, illustrating the phase-locking effect. To improve visibility, the line width $w$ of each curve is modulated based on the correlation $c$ with node $N_1$: $w = 1 + \theta(c) \cdot c \cdot 3$, where $\theta$ is the Heaviside step function. **(d, f, h, j)** Correlation matrices computed from the time series of protein concentrations for each node. **k)** Schematic representation of an alternative (explicit) coupling strategy between oscillators. **l)** The effect of the repression strength between the oscillators ($\beta$) to the phase-locking. In panels (l) and (n), the simulations were run for $10^5$ minutes, with the first 15% treated as transient. Parameters were $a_{1,N2} = 1.1$, $a_{1,N6} = 1.2$, and in (n) [dCas] = 200 molecules. **m)** Schematic illustration of implicit coupling between oscillators mediated by the time-dependent burden. **n)** Phase-locking value versus dCas concentration for various half-saturation constants $K$; lower $K$ indicates stronger burden-mediated coupling.
Promoter strengths were set to 1 for $N_3$ and $N_6$ and to 1.1 for $N_2$ and $N_6$ across all cases, while $a_1$ for $N_1$ and $N_4$ was varied as follows: (a, b) $a_{1,N1} = a_{1,N4} = 1$; (c, d) $a_{1,N1} = 1.2$, $a_{1,N4} = 1$; (e, f) $a_{1,N1} = 1$, $a_{1,N4} = 1.2$; and (g, h) $a_{1,N1} = a_{1,N4} = 1.2$. Simulations were run for $10^5$ minutes, with the first half considered transient. Our previous CRISPRlator model [1] was used for these simulations. Detailed simulation methods are available in the SI under "computer.html."

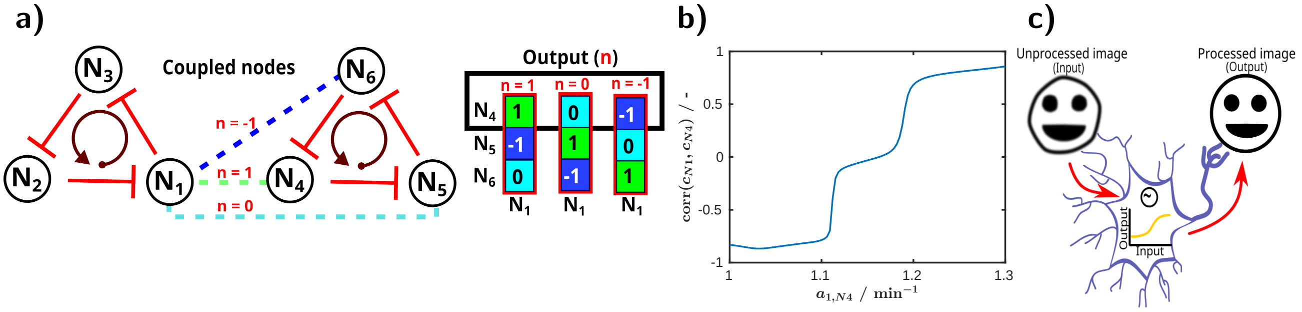

**Figure S21: Oscillator-Based Computation: Phase Locking in Coupled CRISPRlators.** This figure presents an additional analysis of the simulations shown in Figure 6, and illustrates the potential for image processing. **a)** Illustration of ternary output potential beyond binary computation. **b)** Correlation between $N_1$ and $N_4$ as a function of $N_4$ promoter strength. **c)** Schematic illustration highlighting the advantage of using a continuous nonlinear output function over binary signals, with the example of image processing with oscillator-based neurons. For a deeper exploration of image processing techniques like sharpening, we refer readers to [19].

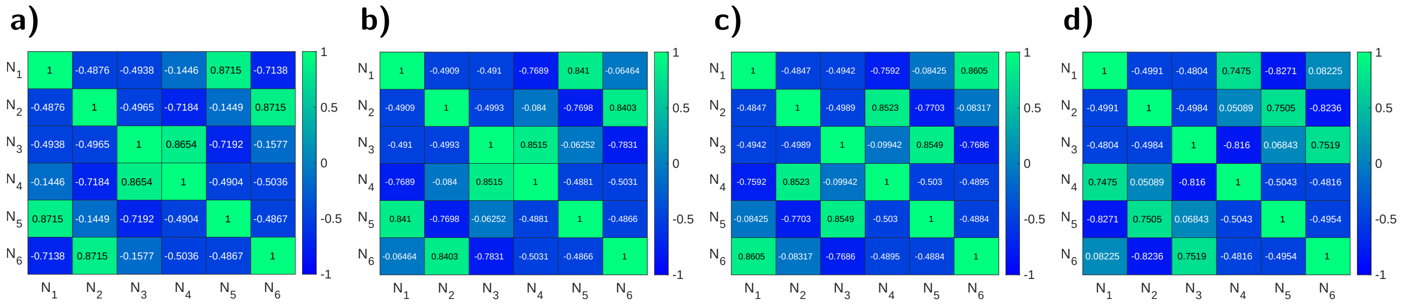

**Figure S22: Oscillator-based computation: demonstration of phase locking with coupled CRISPRlators.** This figure displays the correlation matrices involving all nodes, corresponding to panels d, f, h, and j in Figure 6.

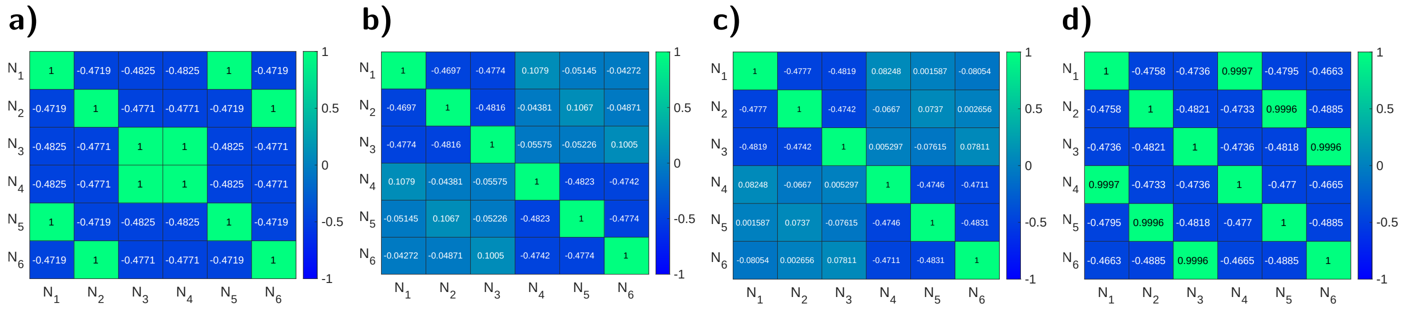

**Figure S23: Oscillator-based computation: demonstration of phase locking with coupled CRISPRlators.** This figure presents the correlation matrices calculated with a dCas concentration of 500 molecules, corresponding to Figure 6 panels d, f, h, and j, respectively.

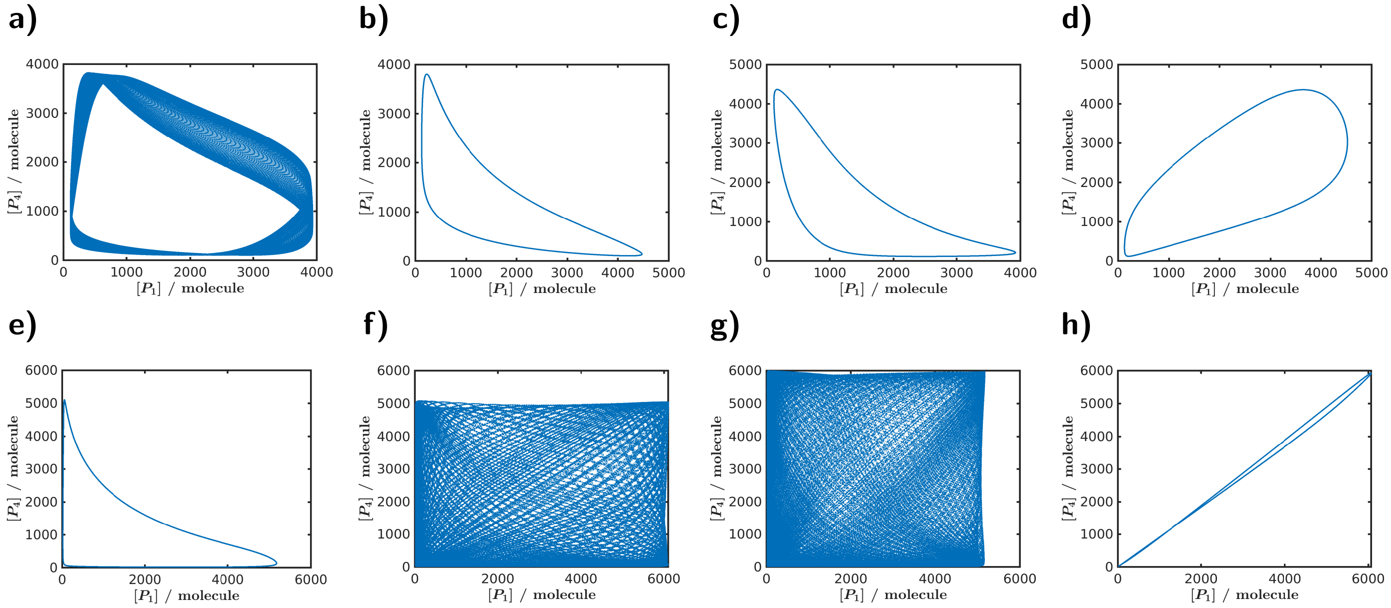

**Figure S24: Oscillator-based computation: Lissajous curves of coupled CRISPRlators in phase space on the $\mathbf{P_{N1}}$-$\mathbf{P_{N2}}$ plane.** The first row (**a**–**d**) corresponds to Figure 6 panels d, f, h, and j, as well as Figure S22 panels a, b, c, and d, respectively. The second row (**e**–**h**) corresponds to Figure S23 panels a, b, c, and d, respectively.

#### 5.1 Stochastic simulations

##### 5.1.1 Relevance for biological oscillator models

Noise is an intrinsic feature of biological systems and must be considered when studying coupled oscillators in practical applications, as it can, for example, disrupt synchronization and thereby alter system behavior. Experimental studies estimated the noise autocorrelation time from protein production rate fluctuations. For intrinsic noise, driven by variations in mRNA copy numbers, the autocorrelation time was found to be $\leq$10 minutes. In contrast, global noise exhibited an autocorrelation time on the order of the cell cycle, $\sim$40 minutes [20, 21]. To capture these processes, we employed Ornstein–Uhlenbeck (OU) noise. Unlike uncorrelated Gaussian white noise, which injects power across all frequencies and can cause unbounded amplitude growth in sustained oscillations, the OU process introduces a finite correlation time $\tau$ that limits the noise bandwidth. Since biological fluctuations typically arise from finite-time processes such as transcription, translation, or molecular transport, OU noise – with its tunable correlation time – offers a more realistic representation of such “colored´ noise.

The dynamics of stochastic biochemical systems can be described by the chemical Langevin equation (CLE). For $N$ molecular species and $M$ reaction channels with stoichiometric change vectors $\boldsymbol{\nu}_j$ and propensity functions $a_j(\mathbf{X})$, the CLE reads

$$\mathrm{d}\mathbf{X}(t) \;=\; \sum_{j=1}^{M} \boldsymbol{\nu}_j \, a_j(\mathbf{X}(t))\,\mathrm{d}t \;+\; \sum_{j=1}^{M} \boldsymbol{\nu}_j \, \sigma \sqrt{a_j(\mathbf{X}(t))}\,\mathrm{d}W_j(t), \tag{S29}$$

where $\sigma$ sets the overall noise strength and $\mathrm{d}W_j(t)$ are independent Wiener processes. While this formulation typically assumes Gaussian white noise, in biological systems it is often more appropriate to consider temporally correlated fluctuations, which we describe using OU noise in the following section.

##### 5.1.2 The Ornstein–Uhlenbeck Process

The dynamics of a one-dimensional OU process are governed by the following stochastic differential equation (SDE):

$$\mathrm{d}X(t) = -\theta X(t)\,\mathrm{d}t + \sigma\,\mathrm{d}W(t), \tag{S30}$$

where $\theta = 1/\tau > 0$ is the mean-reversion rate, $\tau$ is the correlation time, $\sigma > 0$ is the noise intensity, and $W(t)$ is a standard Wiener process.

To derive the exact transition law between discrete time points, we consider the evolution of $X(t)$ over a fixed time step $\Delta t$. The solution to Eq. (S30) can be found using the method of integrating factors. Multiplying both sides by $e^{\theta t}$ yields:

$$e^{\theta t}\mathrm{d}X(t) + \theta e^{\theta t}X(t)\,\mathrm{d}t = \sigma e^{\theta t}\mathrm{d}W(t). \tag{S31}$$

The left-hand side of Eq. (S31) is the stochastic differential of $e^{\theta t}X(t)$, leading to:

$$\mathrm{d}\left(e^{\theta t}X(t)\right) = \sigma e^{\theta t}\mathrm{d}W(t). \tag{S32}$$

Integrating Eq. (S32) from $t$ to $t + \Delta t$ gives:

$$e^{\theta(t+\Delta t)}X(t+\Delta t) - e^{\theta t}X(t) = \sigma \int_t^{t+\Delta t} e^{\theta s}\,\mathrm{d}W(s). \tag{S33}$$

Solving for $X(t + \Delta t)$ and using $\theta = 1/\tau$, we obtain the discrete-time process:

$$X(t+\Delta t) = e^{-\Delta t/\tau}X(t) + \sigma \int_t^{t+\Delta t} e^{-\theta(t+\Delta t-s)}\,\mathrm{d}W(s). \tag{S34}$$

**Statistics of the Integrated Noise Term** The stochastic integral in Eq. (S34) defines the noise increment for the discrete step:

$$\eta_{\Delta t} \equiv \sigma \int_t^{t+\Delta t} e^{-\theta(t+\Delta t-s)}\,\mathrm{d}W(s). \tag{S35}$$

As an Itô integral of a deterministic integrand, $\eta_{\Delta t}$ is a Gaussian random variable with mean zero, $\mathbb{E}[\eta_{\Delta t}] = 0$. Its variance is calculated via the Itô isometry:

$$\begin{aligned}\mathrm{Var}(\eta_{\Delta t}) = \mathbb{E}[\eta_{\Delta t}^2] &= \sigma^2 \int_t^{t+\Delta t} \left(e^{-\theta(t+\Delta t-s)}\right)^2 \mathrm{d}s \\ &= \sigma^2 \int_t^{t+\Delta t} e^{-2\theta(t+\Delta t-s)}\,\mathrm{d}s.\end{aligned} \tag{S36}$$

Making the substitution $u = t + \Delta t - s$, the integral evaluates to:

$$\mathrm{Var}(\eta_{\Delta t}) = \sigma^2 \int_0^{\Delta t} e^{-2\theta u}\,\mathrm{d}u = \frac{\sigma^2}{2\theta}\left(1 - e^{-2\theta\Delta t}\right) = \frac{\sigma^2\tau}{2}\left(1 - e^{-2\Delta t/\tau}\right). \tag{S37}$$

Thus, $\eta_{\Delta t} \sim \mathcal{N}\left(0,\, \frac{\sigma^2\tau}{2}\left(1 - e^{-2\Delta t/\tau}\right)\right)$.

**Exact Discrete-Time Update Formula** From Eqs. (S34) and (S37), the exact state update for any time step $\Delta t$ is given by:

$$X_{n+1} = e^{-\Delta t/\tau} X_n + \sqrt{\frac{\sigma^2 \tau}{2} \left(1 - e^{-2\Delta t/\tau}\right)}\, \xi_n, \tag{S38}$$

where $X_n \equiv X(t_n)$, and $\xi_n \sim \mathcal{N}(0,1)$ is a standard i.i.d. Gaussian random variable. It is often useful to express the update in terms of the stationary (long-term) variance of the process, $\sigma_\infty^2 = \frac{\sigma^2 \tau}{2}$. In these terms, the update rule becomes:

$$X_{n+1} = e^{-\Delta t/\tau} X_n + \sigma_\infty \sqrt{1 - e^{-2\Delta t/\tau}}\, \xi_n. \tag{S39}$$

**Consistency with the Euler–Maruyama Scheme** In the limit of a small time step ($\Delta t \ll \tau$), the exact discretization reduces to the Euler–Maruyama scheme. Using the first-order approximations $e^{-\Delta t/\tau} \approx 1 - \Delta t/\tau$ and $e^{-2\Delta t/\tau} \approx 1 - 2\Delta t/\tau$ in Eq. (S38) yields:

$$X_{n+1} \approx \left(1 - \frac{\Delta t}{\tau}\right) X_n + \sigma \sqrt{\Delta t}\, \xi_n, \tag{S40}$$

which is precisely the Euler–Maruyama discretization of the SDE in Eq. (S30). This confirms the consistency of the exact discretization for infinitesimal time steps.

###### 5.1.3 Methods

The stochastic simulations were carried out for the CLE using the Euler–Maruyama method with the OU noise using the exact discrete-time formula (Equation S39), with the solver implemented in C++ as a `mex` function to interface with MATLAB. Each simulation was run for $10^5$ minutes with a step size of 0.01 minute, with $\tau = 10$ minutes correlation time, and the first 15% of the trajectory was discarded as transient dynamics.

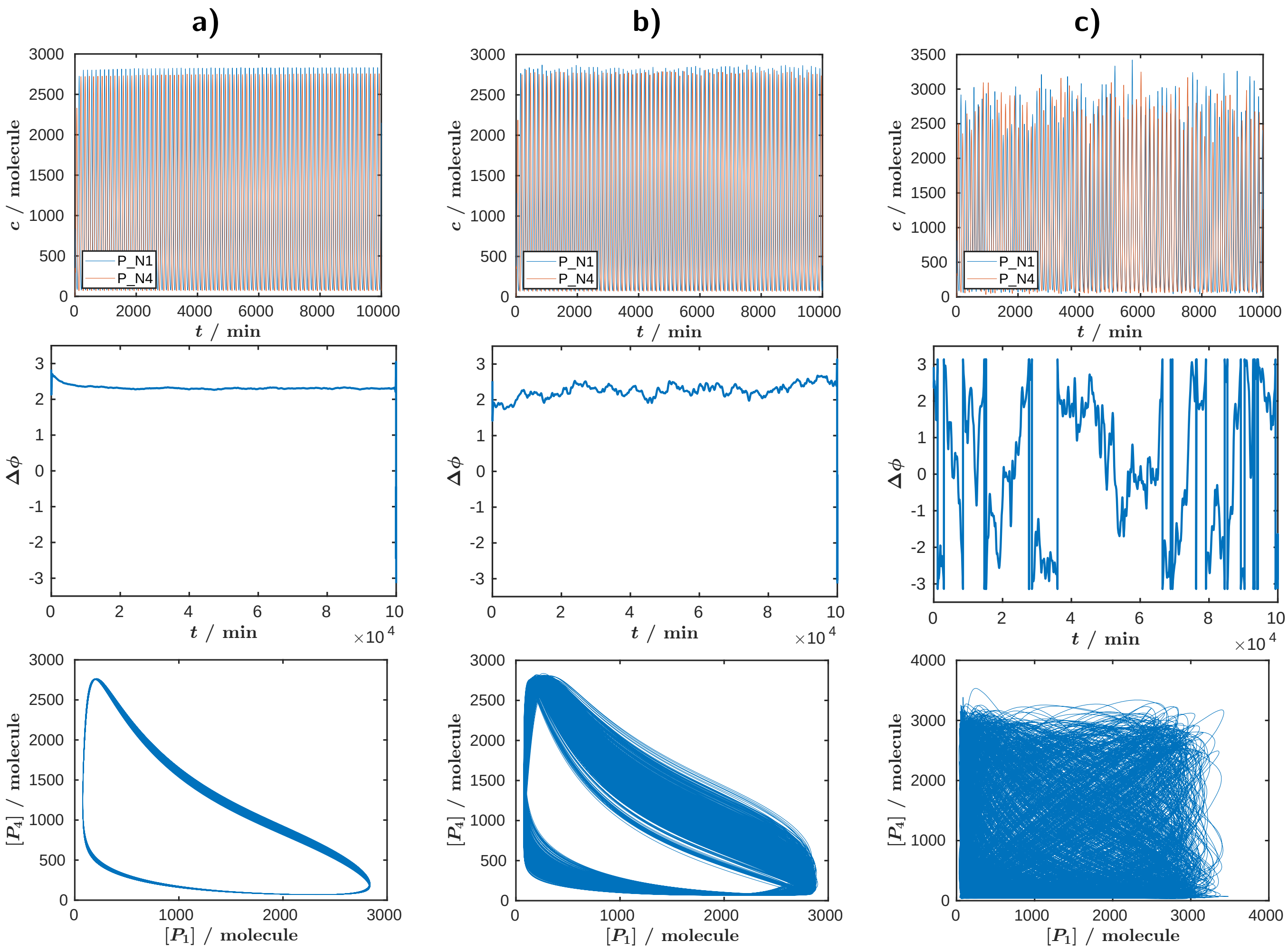

**Figure S25: Example simulations with varying noise strength.** Top row: trajectories over the first $10^4$ minutes of the $10^5$ minutes long simulations. Middle row: instantaneous phase difference (between $P_{N1}$ and $P_{N4}$) as a function of time. Bottom row: stochastic Lissajous curves. The noise strength $\sigma_\infty$ was set to $10^{-4}$, $10^{-3}$, and $10^{-2}$ in panels (a), (b), and (c), respectively. These simulations correspond to Figure 6l, with the half-saturation constant for the burden fixed at $K = 10^4$ molecules. For the Lissajous curves, the first 15% of each simulation was omitted as transient behavior. See Section 5.1 for details of the stochastic simulations.

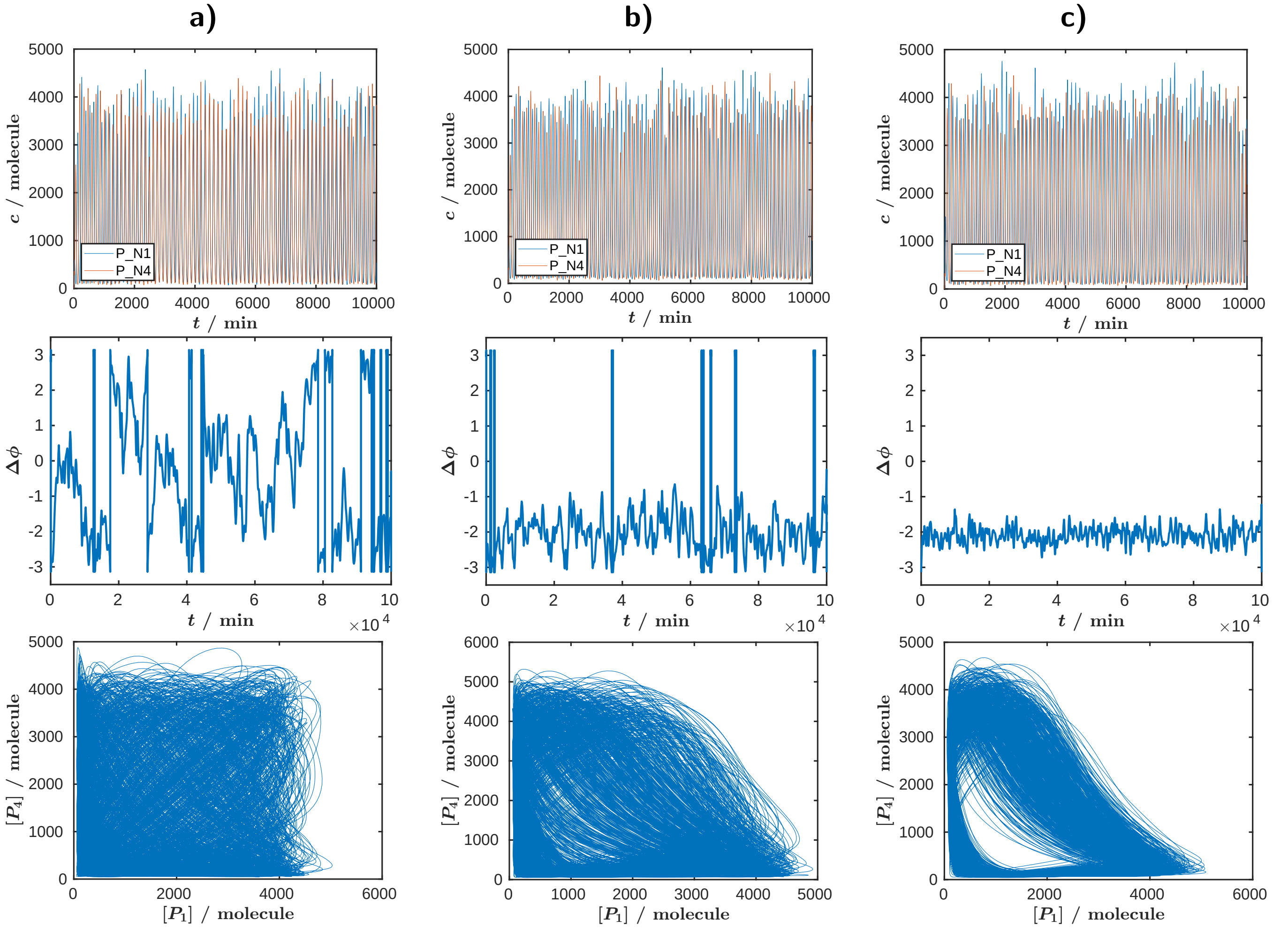

**Figure S26: Example simulations with varying repression strength between the oscillators (Figure 6k,l).** Top row: trajectories over the first $10^4$ minutes of the $10^5$ minutes long simulations. . Middle row: instantaneous phase difference (between $P_{N1}$ and $P_{N4}$) as a function of time. Bottom row: stochastic Lissajous curves. The repression strength was set to 0, 0.01, and 0.1 times of the regular repression strength in panels (a), (b), and (c), respectively, while the noise strength $\sigma_\infty$ was set to $10^{-2}$. For the Lissajous curves, the first 15% of each simulation was omitted as transient behavior. See Section 5.1 for details of the stochastic simulations.
